# Estimation of non-additive genetic variance in human complex traits from a large sample of unrelated individuals

**DOI:** 10.1101/2020.11.09.375501

**Authors:** Valentin Hivert, Julia Sidorenko, Florian Rohart, Michael E Goddard, Jian Yang, Naomi R Wray, Loic Yengo, Peter M Visscher

## Abstract

Non-additive genetic variance for complex traits is traditionally estimated from data on relatives. It is notoriously difficult to estimate without bias in non-laboratory species, including humans, because of possible confounding with environmental covariance among relatives. In principle, non-additive variance attributable to common DNA variants can be estimated from a random sample of unrelated individuals with genome-wide SNP data. Here, we jointly estimate the proportion of variance explained by additive 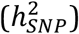, dominance 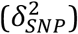 and additive-by-additive 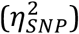 genetic variance in a single analysis model. We first show by simulations that our model leads to unbiased estimates and provide new theory to predict standard errors estimated using either least squares or maximum likelihood. We then apply the model to 70 complex traits using 254,679 unrelated individuals from the UK Biobank and 1.1M genotyped and imputed SNPs. We found strong evidence for additive variance (average across traits 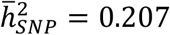. In contrast, the average estimate of 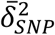 across traits was 0.001, implying negligible dominance variance at causal variants tagged by common SNPs. The average epistatic variance 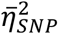 across the traits was 0.058, not significantly different from zero because of the large sampling variance. Our results provide new evidence that genetic variance for complex traits is predominantly additive, and that sample sizes of many millions of unrelated individuals are needed to estimate epistatic variance with sufficient precision.

## Introduction

The total genetic variance of a trait can be partitioned into additive, dominance and epistatic variance components^1–5^. The role of non-additive genetic variation (dominance and epistatic) in complex traits in human populations remains elusive, because it is difficult to estimate and also because theory predicts it to be small relative to additive variance. Traditionally, non-additive genetic variance has been estimated from pedigree or twin studies, by contrasting the phenotypic covariance for different kinds of relatives. Non-additive variance leads to closer relatives being more similar than expected from additive genetic variance. However, shared environmental effects may also be expected to be strong among closer relatives, and disentangling these and other sources of familial resemblance remains challenging.

The amount of non-additive genetic variance disproportionally depends on the allele frequencies at causal variants as compared to additive variance, and in general it is lower when such frequency (and locus heterozygosity) is low. Non-additive genetic variance can be estimated in model species, where both the environment and allele frequencies can be controlled^6^. Theory and empirical evidence suggest that genetic variation in complex traits is mainly additive^7,8^ and that higher-order epistatic interactions, if present, are still mostly expected to contribute to additive genetic variance^8^. For highly polygenic traits with evidence of directional dominance (also known as inbreeding depression), the expected dominance variance is also very small^9^.

Variance components can also be estimated from genome wide SNP genotypes. In humans, Zhu et al.^10^ extended the model of Yang et al.^11^ to estimate the proportion of phenotypic variance explained by both additive 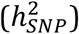 and dominance variance 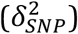 from common SNPs in a sample of unrelated individuals using restricted maximum likelihood (REML). Across 79 quantitative traits in a sample of 6,715 individuals, they reported an average dominance variance of 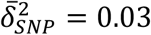, approximately one-fifth of their estimated average narrow sense SNP-based heritability 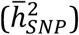, with a large standard error 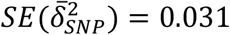 (computed using the reported standard errors of Zhu et al.^10^ and neglecting the covariance between estimates). Therefore, these results are consistent with a small to negligible contribution of dominance variance across these traits.

Epistatic variance attributable to common DNA variants 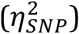 can, in principle, also be estimated from a random sample of unrelated individuals with genome-wide SNP data. However, pairwise coefficients for epistatic variance between unrelated individuals are very small, and *a priori* it is expected that non-additive variance contributes less than additive variance. How large sample sizes need to be to reliably detect epistatic variance is currently unknown. Therefore, having a theoretical expectation of the sampling variance of estimators of 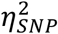 would allow the quantification of statistical power and thus the required sample size. Moreover, theory predicts that imperfect tagging of causal variants by SNPs leads to a larger reduction in SNP-based estimates of non-additive genetic variance as compared to SNP-based estimates of additive genetic variance. For example, if a causal variant is tagged by a SNP with an linkage disequilibrium (LD) squared correlation r^2^, then estimates of additive variance are expected to be proportional to r^2^, whereas it is r^4^ for dominance and additive-by-additive variance^12^.

Our study focuses on additive-by-additive epistatic variance as opposed to that generated by higher-order interactions such as dominance-by-additive or dominance-by-dominance components. We derive the theoretical sampling variance of ordinary least squares using Haseman-Elston (HE) regression, and Restricted maximum likelihood (REML) estimators of 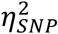 in a sample of unrelated individuals. We jointly estimate the genome-wide additive, dominance, and additive-by-additive SNP-based genetic variances of 70 human complex traits using a large sample of unrelated individuals (*N* = 254,679) from the UK Biobank^13^ (UKB).

## Results

### Overview of the method

Zhu et al.^10^ developed a linear mixed model (LMM) where the total genetic variance of a trait attributable to SNPs is partitioned into additive and dominance variance and where the effects of all genotyped or imputed SNPs are fitted together as random effects through the use of additive and dominance genomic relationship matrices (GRM). The different variance components of this multiple-GRM model were then estimated using REML. Here, we extend this model by partitioning the total genetic variance of a trait that is attributable to SNPs into additive, dominance and additive-by-additive variance components. That can be mathematically written as:

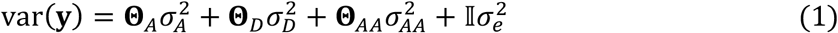

with **y** a vector of phenotypic values for N diploid individuals and 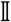 an *N* × *N* identity matrix. The three components of genetic variance 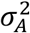, 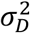 and 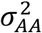 are respectively the additive, dominance and additive-by-additive variance explained by SNPs, and 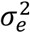 is the residual variance. **Θ**_***A***_ = *G* and **Θ**_*D*_ are the additive and dominance GRMs computed from L SNPs as previously described in Yang et al.^11^ and Zhu et al.^10^ (see **Methods** for details), and **Θ**_***AA***_ is the additive-by-additive genomic relatedness matrix, defined as:

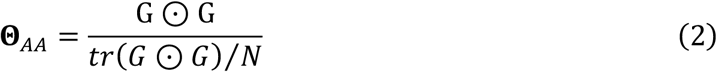

with G ☉ G the Hadamard product (i.e. coefficient-wise matrix product) of the additive GRM with itself, and *tr*(*G* ☉ *G*) the trace of the matrix. The standardization by the average of the diagonal elements ensures that the mean of the diagonal elements of **Θ**_***AA***_ is approximately one as for **Θ**_*A*_ and **Θ**_*D*_ (**Supplementary Table 1**), leading to estimates of genetic variances on the same scale as the residual variance^14,15^.

Under Hardy-Weinberg equilibrium, the model defined by **Equation (1)** is theoretically orthogonal between the additive and dominance components (see **Methods** and Zhu et al.^10^). By orthogonality, we mean that the estimates of one of the genetic variance components when fitted as the only genetic component would be unbiased even in the presence of the effects from the other components. Moreover, in a sample of unrelated individuals from an outbred population, the off-diagonal elements of Θ_*A*_ are expected to be 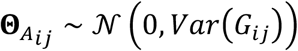, leading to an expected covariance 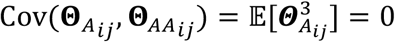. The latter property guarantees orthogonality between the additive and additive-by-additive component. Orthogonality between the dominance and additive-by-additive components is more complicated to prove and was investigated empirically.

**Equation (1)** defines a typical LMM, which variance components 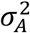, 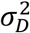, 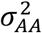 and 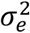) can be estimated jointly or separately using REML^16^ as well as by HE regression^17–19^. The proportion of phenotypic variance explained by additive 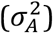, dominance 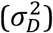 and additive-by-additive 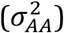 variance at all SNPs are defined as 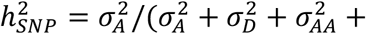 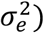, 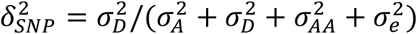 and 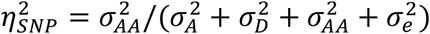. We define the SNP-based broad sense heritability 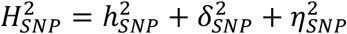. Details about the statistical model and the computation of the different GRMs are provided in the **Methods** section.

### Unbiased estimates when all GRMs are fitted simultaneously

We use simulation to quantify empirically the bias and precision of the HE and REML estimators of genetic variances, as well as to assess consistency with theory. We first validate our model using simulations from 254,679 unrelated participants of the UKB with 1.1M autosomal genotyped and imputed SNPs from the HapMap 3 (HM3) panel^20^ with minor allele frequency (MAF) ≥ 1%. In our simulations, we sampled putative causal variants from a pool of 100,000 pre-defined HM3 SNPs (**Methods**). We used the same pool of causal variants across all simulation replicates from which we randomly sampled 1000 causal variants for each replicate. The phenotypes were simulated using **Equation (5)**(in **Methods**) where the additive, dominance, and additive-by-additive effects were generated from a standard normal distribution and adjusted to the expected variance of the additive, dominance and additive-by-additive genome-wide effects (simulated 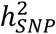, 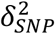 and 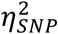). The residuals were then generated from a normal distribution with mean 0 and variance 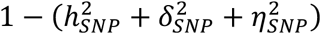. The genotype data were used to calculate the three GRMs *Θ*_*A*_, *Θ*_*D*_ and Θ_*AA*_ as defined above (summary statistics for each GRM are provided in **Supplementary Table 1 and 2**).

We estimated 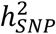, 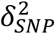 and 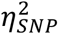 either jointly (hereafter referred to as ADAA model) or by including their corresponding GRM one at a time in the model. Variance components were then estimated using REML and HE regression, which are both implemented in the GCTA^16^ software. The full dataset of 254,679 individuals was analyzed using HE regression with pairwise phenotypic cross-products. However, because of its heavy computational burden, REML estimates were derived from 8 sub-datasets, each of ≃ 32K individuals. We meta-analyzed estimates from these 8 sub-datasets using inverse-variance weighting (IVW) (**Methods**), recognizing that the standard errors of these meta-analysed estimates are expected to be 2.8 fold higher than if analysis of a single combined dataset had been possible for additive and dominance variance, and 1.3 fold higher for additive-by-additive variance. Unless stated otherwise, simulations results are shown for analyses including the causal variants.

From REML analysis, the model shows good orthogonal properties (fitting all the three GRMs together or only one at a time does not change the estimates) for 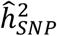and 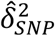. However, we found a significant deviation from regressing 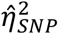 estimates from the ADAA model and that from a univariate model fitting Θ_*AA*_ only (intercept = 0.65, see **Figure 1-A**). The same conclusion applies to HE analysis where 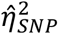 from the univariate model show an even stronger deviation (intercept = 1.20) from the joint estimates (**Figure 1-B**), as compared to REML. A lack of orthogonality is expected when using linked markers as orthogonal estimates of epistatic variance are not possible^21^. However, we observed a lack of orthogonality in simulations using simulated unlinked markers and unrelated individuals (**Supplementary Figure 1**). Although this observation was initially surprising, we derived theoretically why this occurs, and reflects induced collinearity between the GRM because allele frequencies are estimated with error from finite sample sizes (**Supplementary Note 1** and **Supplementary Figure 9**). The main and practical conclusion from these simulations is that *Θ*_*A*_ and *Θ*_*D*_, the additive and dominance GRMs, should always be fitted when estimating additive-by-additive variance using Θ_*AA*_.

**Figure 1:**
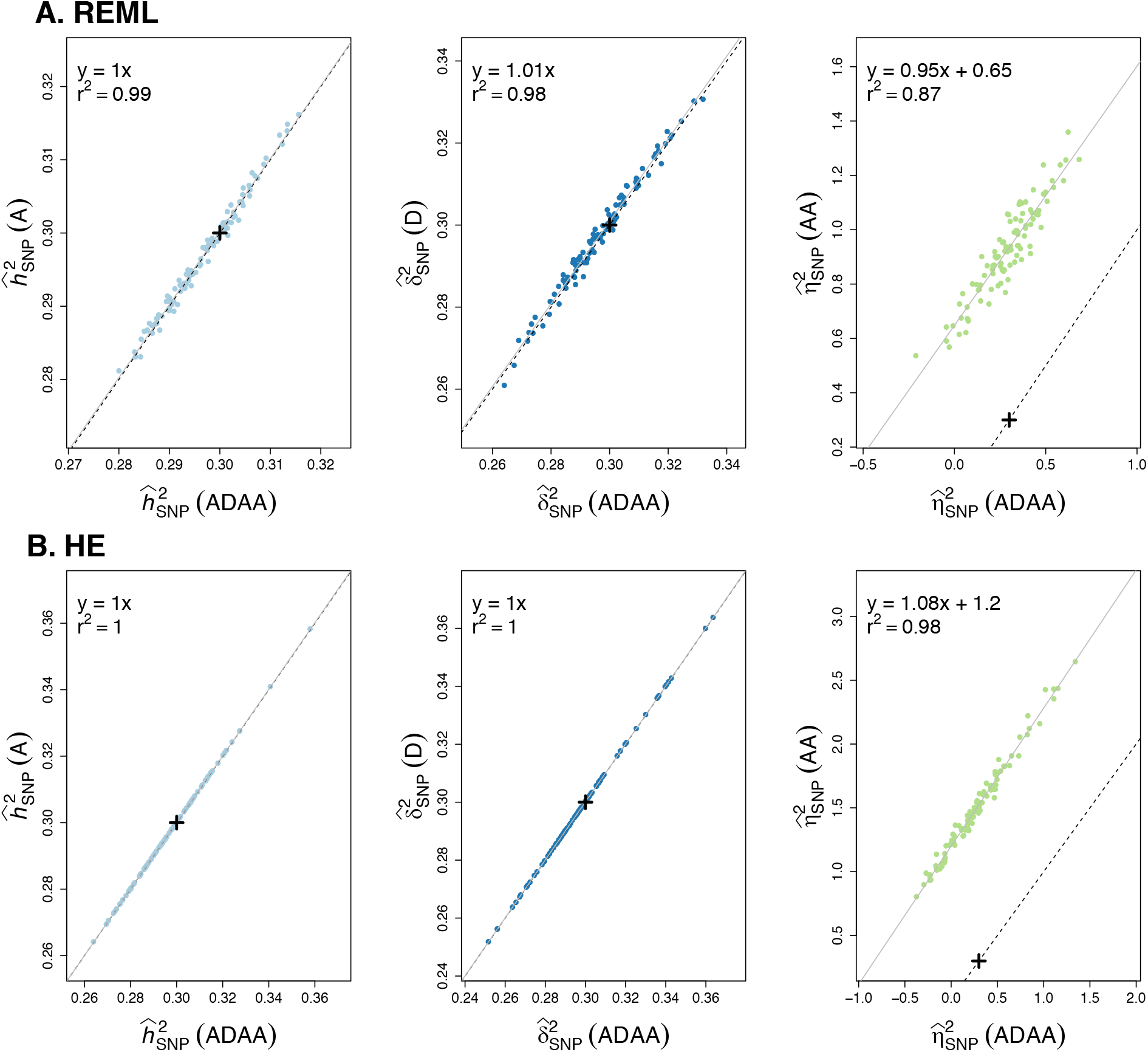
REML and HE estimates from simulations based on observed genotypes of UKB participants (simulated 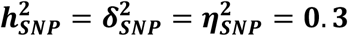). We compared (**A**) IVW-REML (meta-analysis of 8 sub-datasets of 32K individuals) and (**B**) HE estimates (analysis of 254,679 individuals) for 100 replicates of simulations including the pool of causal variants when we jointly estimate the different variance components (ADAA) or only one at a time for 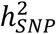, 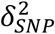 and 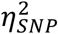. Simulated values are depicted by a black cross, the black dashed line depicts the Y=X line while the solid grey line depicts the linear regression between the corresponding ADAA and single component model. The squared correlation r^2^ is close to 1 for all the variance components and η^2^ is the only one showing a large intercept.

After assessing the orthogonality properties of the model, we evaluated its performance with respect to loss of information due to tagging and its sampling variance. Consistent with Yang et al.^22^, we define unbiasedness as when the average estimates across simulation replicates is not statistically different from the true expected value (t-test *P*-value>0.05). **Figure 2** shows that REML analysis performed with all observed HM3 SNPs including causal variants yields nearly unbiased estimates of the three variance components 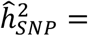 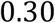, 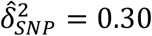,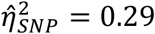. A slight downward bias is expected when dealing with missing genotypes that are imputed during the simulation process. Using simulation of unrelated individuals and unlinked markers, we can show that 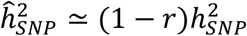 with r the missing genotype rate (**Supplementary Figure 4**). When the pool of causal variants was excluded from the analysis 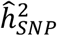 and 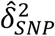 both showed a downward bias with respective mean estimates of 0.28 and 0.26 whereas 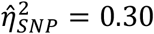 did not appear deflated. However, we lacked power to detect a small bias because of the large sampling variance of 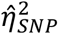. To verify that the biases for 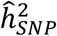 and 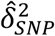 are consistent with loss of tagging, we estimated the adjusted multiple correlation *R*^2^ computed for the first 10K causal SNPs with nearby SNPs in 1Mb windows (**Methods**). We found values of 0.96 and 0.86, for additive and dominance effects, respectively, leading to expected values 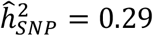 and 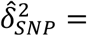 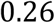, consistent with the estimated variance components. These results show that despite incomplete LD we are still able to detect additive and dominance variation due to common causal variants in the UKB data. Results from HE analysis imply the same conclusion (**Figure 2**). Finally, we found the standard deviation (SD) of 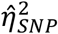 across simulation replicates to be one order of magnitude larger than that of 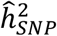 and 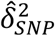. More specifically, 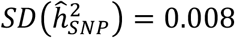, 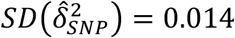 and 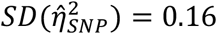, in agreement with theoretical expectations presented below.

**Figure 2:**
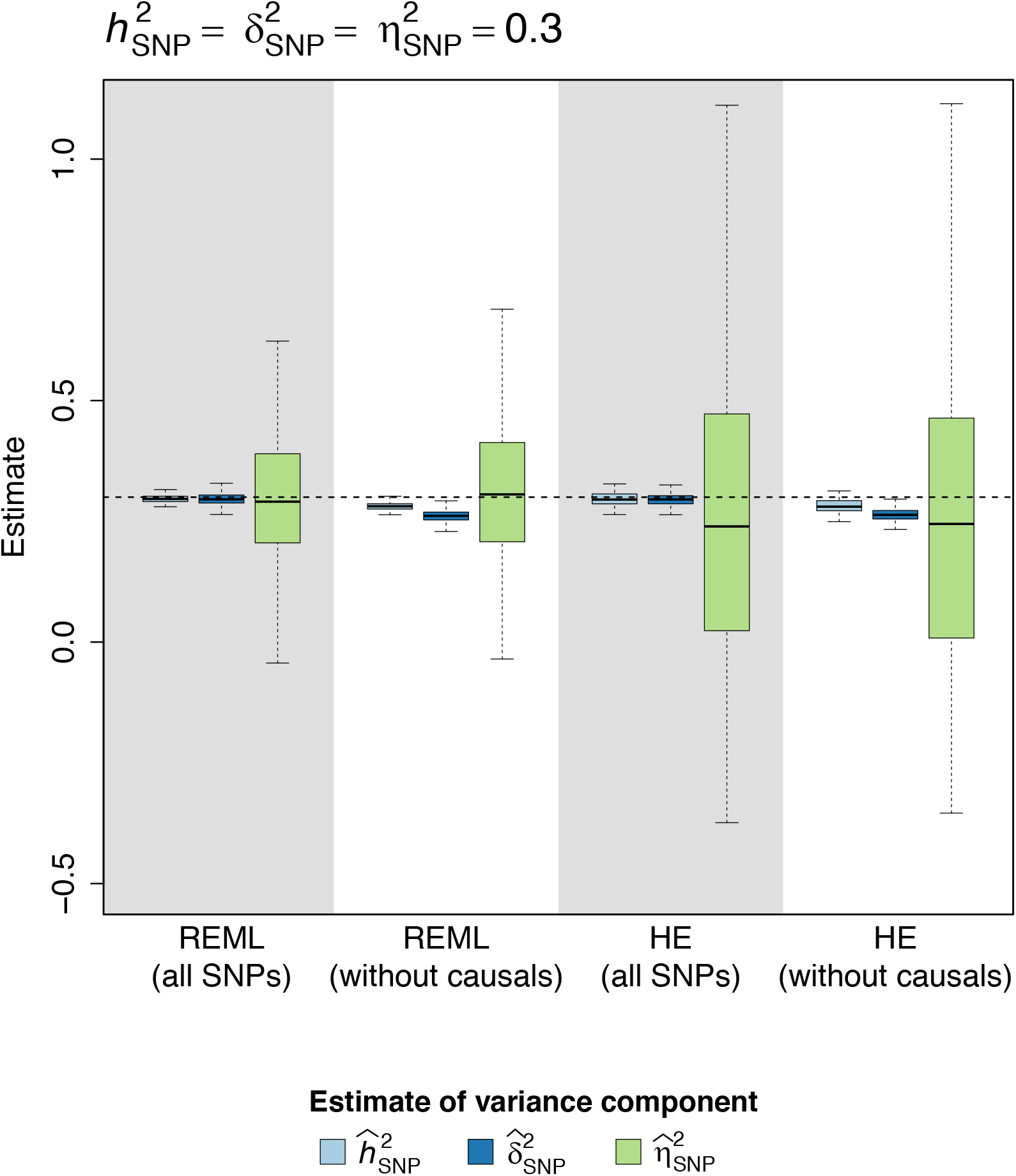
Distributions of REML and HE estimates from simulations based on observed genotypes of 254,679 UKB participants (simulated 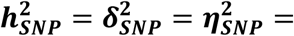 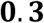). IVW-REML (analysis of 8 sub-datasets of 32K individuals) and HE estimates (analysis of 254,679 individuals) for 100 replicates of simulations are shown for the three different variance components including all variants (all SNPs) or excluding the causal variants (without causal). The black dashed line indicates the simulated value of 0.3.

### Standard error of the estimate of additive-by-additive variance

We next sought to derive the theoretical sampling variance of estimators of 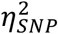 Assuming a sample of *N* unrelated individuals from an outbred population, the diagonal and off-diagonal elements of the G matrix are respectively 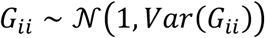 and 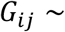 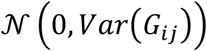. Using results from Visscher et al.^23^ and Visscher and Goddard^24^, we derived different sampling variance for HE regression using phenotypic cross-products and REML (**Supplementary Note 2**),

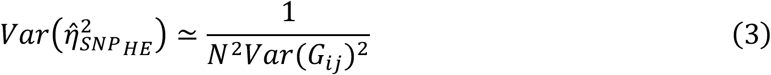

and

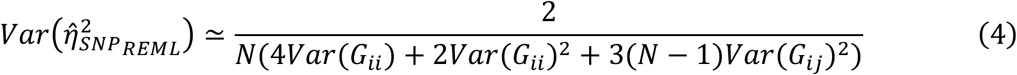

This result implies that statistical power to detect 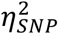 is substantially larger when using REML as compared to HE. Note that the sampling variance 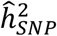 of and 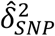 under REML or HE regression are approximately the same. For an infinite sample size, 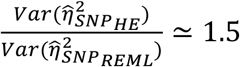 (**Supplementary Note 2**), while this ratio is in fact much larger for finite sample sizes (**Supplementary Figure 2**).

From simulations of 35K unrelated individuals with simulated unlinked markers, we found approximations from **Equations (3) and (4)** to be accurate. For REML analyses across 100 replicates, we observed 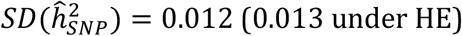, 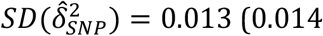 under under HE) and 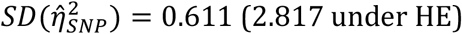, as compare to theoretical standard errors 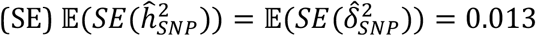 (under REML and HE), and 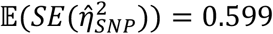 under REML (2.858 under HE). We then used simulations based on actual genotypes from UKB participants to assess the accuracy of our theoretical expectation on real data. For each replicate, we quantified the empirical SE of estimates of variance components as the SD across 8 sub-datasets of 32K individuals. The observed standard deviations were then averaged over replicates and compared to the theoretical expectations. Overall, we found both our theoretical expectations to be accurate (**Table 1**). We noticed a very small absolute bias downward for the theoretical SE in the order of 10^−3^ for 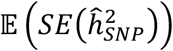 and 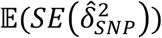, and upward in the order of 10^−2^ for 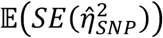, both under REML and HE.

**Table 1:** Observed standard deviations (SD) from the analysis of simulations based on observed genotypes of 254,679 UKB participants (si mulated 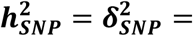 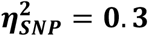), as well as theoretical and reported standard errors (SE) by GCTA.

| Analysis | Variance component | Observed SD | Theoretical SE | reported SE |
| --- | --- | --- | --- | --- |
| $REML_{32K}$ | $\widehat{h^2}$ | 0.011 | 0.010 | 0.012 |
| | $\widehat{\delta^2}$ | 0.015 | 0.014 | 0.016 |
| | $\widehat{\eta^2}$ | 0.37 | 0.399 | 0.393 |
| $HE_{32K}$ | $\widehat{h^2}$ | $1.4 \times 10^{-2}$ | $9.9 \times 10^{-3}$ | $9.9 \times 10^{-3} (1.8 \times 10^{-2})$ |
| | $\widehat{\delta^2}$ | $1.8 \times 10^{-2}$ | $1.4 \times 10^{-2}$ | $1.5 \times 10^{-2} (2.2 \times 10^{-2})$ |
| | $\widehat{\eta^2}$ | 1.45 | 1.57 | 1.55 (2.19) |

We performed 100 replicates of simulations based on observed genotypes of 254,679 UKB participants. In order to assess the accuracy of the theoretical and reported SE, 8 sub-datasets of 32K individuals were analyzed for each replicate both with REML ( $REML_{32K}$ ) and HE ( $HE_{32K}$ ). The observed SD have been computed within a replicate and averaged across them for each variance component. Reported SE by GCTA were averaged within and across replicates and the format for HE analysis is $SE_{OLS}(SE_{jackknife})$ .

We also re-assessed the validity of the SE reported by GCTA and found it to be accurate for REML (**Table 1)**. When performing HE analysis, GCTA reports two estimates of SE based on OLS (*SE*_*OLS*_) or jackknife (*SE*_*jack*_). Comparing the observed SD to the reported SE from GCTA, we found *SE*_*OLS*_ to be slightly biased downward for 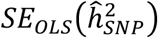 and 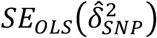 and upward for 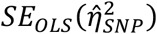 as observed with the theoretical SE. However, the reported *SE*_*jack*_ by GCTA is always more conservative. Therefore, we chose to use the latter and the corresponding *P*-values for the real phenotype analysis.

Finally, using our theoretical expectations, we computed the expected power to detect a significant additive-by-additive effect from our UKB data. The sampling variance of 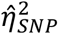 is expected to be very large under HE for small sample sizes, when it is much smaller under REML (**Supplementary Figure 2**), leading to a substantially larger power (**Figure 3**). However, the increase of sample size reduces the gap between the two approaches and we expect 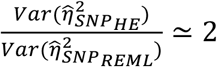 for a sample size of 1M unrelated individuals (**Supplementary Figure 2**). We show that even if we were to be able to analyze 254K individuals jointly with REML, our power (at *α* = 5%) to detect an additive-by-additive effect of 0.2 (which is unlikely in real data) would only be ≃0.45 under REML and ≃0.17 under HE. Under the REML IVW meta-analysis, the expected standard error becomes 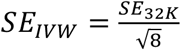, where *SE*_32*K*_ is the expected SE for a sample of 32K individuals. Hence, our expected power under REML slightly decreases because of the meta-analysis strategy (**Supplementary Figure 3**) and we expect a power of ≃0.29 (*α* = 0.05) for 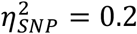. Comparing the expected SE under the IVW-REML and the HE analysis of 254K individuals, we expect slightly larger 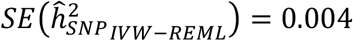 and 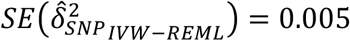 as compared to 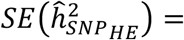 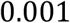 and 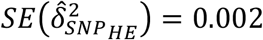. However, the meta-analysis under REML is expected to produce a smaller 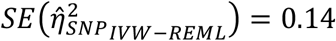 as compared to 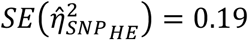.

**Figure 3:**
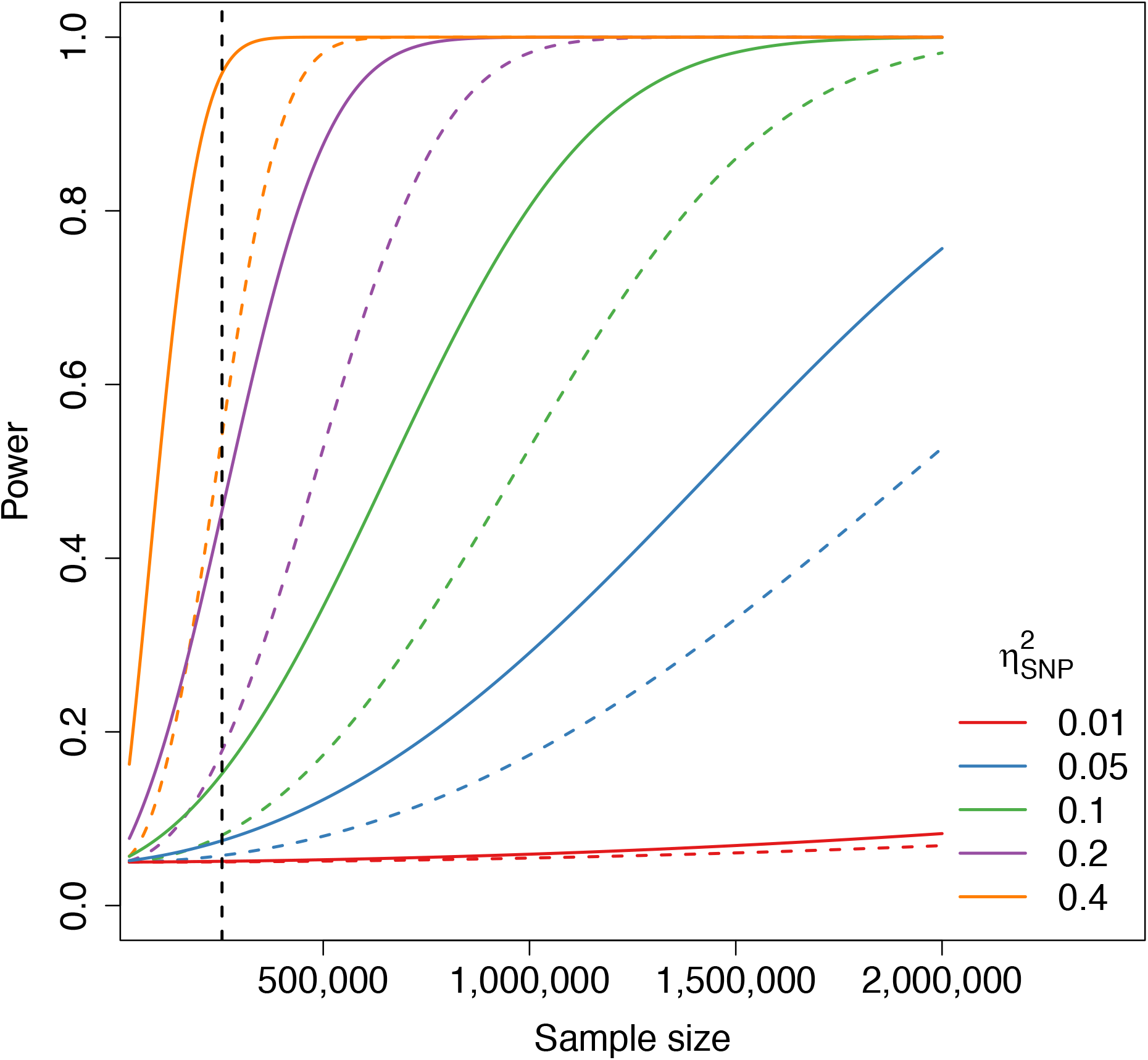
Statistical power to detect additive-by-additive variance as a function of sample size for REML (solid lines) and HE regression (dashed lines) on additive-by-additive GRMs computed on UKB data. Results are shown for an additive-by-additive heritability equal to 0.01, 0.05, 0. 1, 0.2 and 0.4. The current sample size of 254,679 unrelated individuals is depicted by the vertical black dashed line.

In summary, our simulations and analytical results demonstrate that reliable inference of non-additive variance components is achievable using REML and HE regression in large samples, and therefore that these methods can be applied to real data.

### Estimation of non-additive genetic variation in human complex traits

We estimated 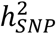, 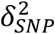 and 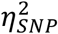 for 70 quantitative traits of the UKB (**Methods**), with an average number of phenotyped individuals ≃ 200,550 (range: 36,990 to 251,805). For REML analysis, IVW meta-analysis was applied when *N* > 60*K*. Otherwise, the entire set of individuals was analysed (**Methods**). Estimates of 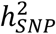, 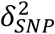 and 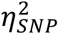 using REML and HE are reported in **Supplementary Tables 4 and 5** and the distributions of REML and HE estimates across traits are shown in **Figure 4**. The mean estimate of 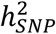 across traits was 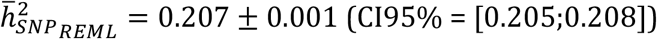 for REML and slightly lower for HE with 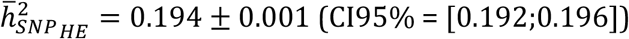. The average estimate of 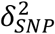 across traits was 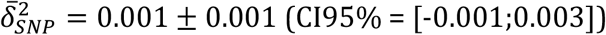 for both REML and HE. This result is in agreement with theory suggesting than 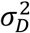 is much smaller than 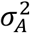. Finally, we found a mean estimate of 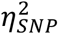 equal to 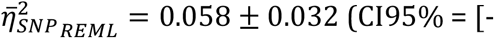 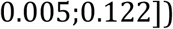 for REML and 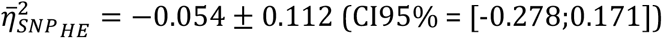 for HE. Estimates are consistent between REML and HE analysis for 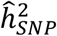 and 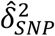 (squared correlations r^2^=0.97 and 0.43 respectively), but the two methods show poor agreement for 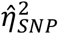 (squared correlation r^2^=0.04, see **Supplementary Figure 5**), consistent with their high standard errors.

**Figure 4:**
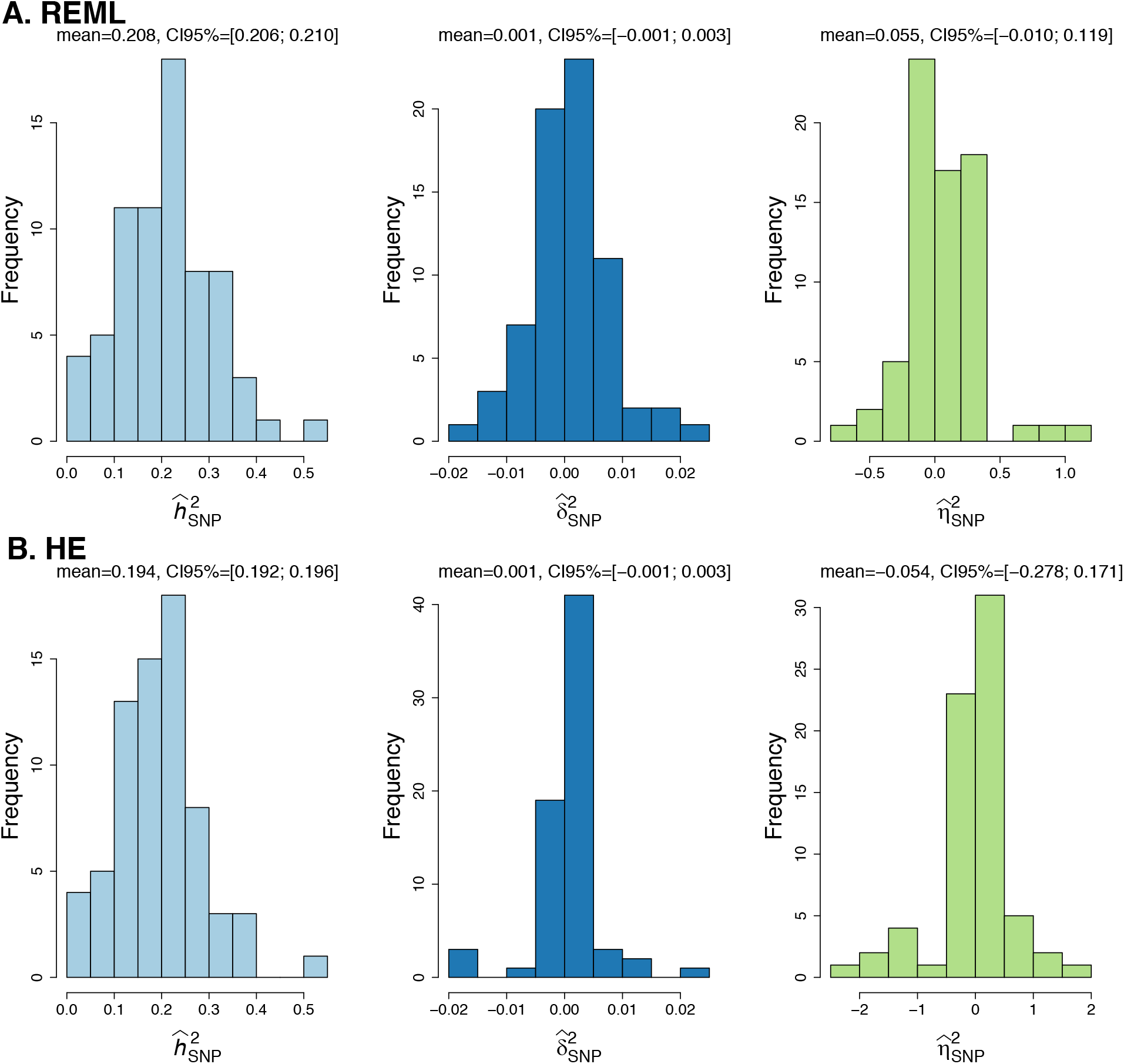
Distributions of the (A) REML and (B) HE estimates of SNP-based 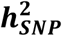 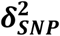 and 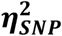 for 70 continuous traits in the UK Biobank. For each distribution of variance components estimates, we indicate the mean estimate as well as the 95% confidence interval (CI95%).

From the 70 traits analysed, we estimated an effective number of 43 traits (**Methods**) and test for 3 variance components, however, because we ascertained traits with significant additive variance (**Methods**), we only included two variance components for multiple testing correction. Therefore, our *P*-value threshold for declaring statistical significance at an experiment-wise error rate *α* = 0.05 is 0.05/(43 × 2) = 5.8e-4. After multiple-testing correction, urate concentration (N = 238,773) was the only trait showing a small but significant dominance variance 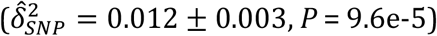 under HE analysis. However, significance was not obtained using REML 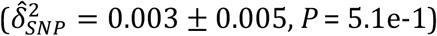.

We did not find significant evidence of epistatic variance. However, we unexpectedly found two traits with an apparent negative 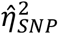 but significant before multiple-testing correction under REML, Bone mineral density (BMD) 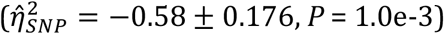 and Corneal resistance factor 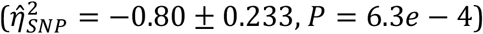. This unusual result implies that close relatives would be disproportionally less phenotypically similar than distant relatives. We therefore quantified phenotypic covariance in family data in the UKB as a function of relatedness but found no evidence for “phenotypic repulsion”^25^ (**Supplementary Figure 6**). All these estimates of non-additive genetic variation are implausibly large, so the most parsimonious explanation is the large sampling variance.

## Discussion

The role of non-additive genetic variance in human complex traits has been a topic of much discussion and debate^7,26–30^. In this study, we jointly estimate the proportions of phenotypic variance of 70 traits that is explained by additive, dominance and additive-by-additive genetic variation tagged by common SNPs, in a large sample of 254,679 unrelated individuals from the UKB. Using common variants, we found no evidence of significant non-additive variance (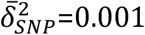 and 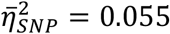) across traits while we confirm the evidence for additive variance 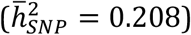. We also derived theoretical standard errors for REML and HE regression estimators of additive-by-additive variance and validated our theory through extensive simulations. Our theoretical and empirical results suggest that REML should be preferred over HE regression for the same sample size, as the former estimator leads to a substantially larger statistical power.

A lack of dominance variance is expected from theory^7^ and for traits that are polygenic and subjected to inbreeding depression.^9^ However, most of our 70 traits do not show evidence of inbreeding depression. Nevertheless, the argument of high degree of polygenicity could be enough to expect a lack of dominance variance too^31,32^. Indeed and as quoted by Crow^31^ for continuous quantitative traits, “in general, the smaller the effects, the more nearly additive they are” (see **Supplementary Note 3**). In a large sample, our average dominance variance estimated across traits 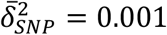, is much lower than the 0.03 previously reported by Zhu et al.^10^ across 79 traits, although the two estimates are not significantly different from each other. Therefore our findings are consistent with that from Zhu et al.^10^ as they confirm a negligible contribution of dominance variance for causal variants that are tagged by common SNPs. By extrapolation, our results lead to the conclusion that dominance variance is likely to contribute very little to the broad sense heritability of human complex traits. Urate concentration was the only trait showing a significant dominance variance in our HE analysis (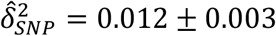, *P* = 9.6e-5), but the estimate was lower and non-significant in our REML analysis (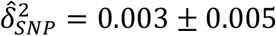, *P* = 5.1e-1). Similarly, we observed that 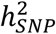 estimates for urate concentration were higher using HE compared to REML (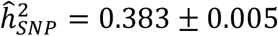 under HE and 0.304 ± 0.004 under REML). If REML and HE estimates are expected to converge under a classical polygenic model, the genetic architecture of a trait can bias the estimates of the two methods in different ways^33–35^. Previous studies^36,37^ have suggested that urate concentration is a trait on the low spectrum of polygenicity and controlled by three main genes and two large effect QTLs on chromosome 4. Moreover, urate concentration also displays sex-specific effects and heritability^38^. Analysis performed on each sex separately (with a 3 fold IVW meta-analysis for REML) confirmed a larger estimated heritability in females 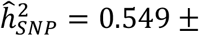 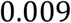 under HE and 0.354 ± 0.005 under REML) than in males 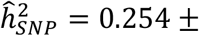 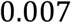 under HE and 0.237 ± 0.006 under REML), and a significant dominance variance for females only under HE analysis (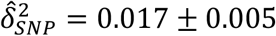, *P* = 1.4e-3 for females, and 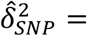 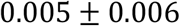 ±, *P* = 3.9e-1 for males). Importantly, the significant dominance variance estimate detected in females with HE vanishes when excluding SNPs on chromosome 4 from our analysis (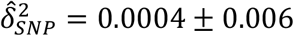, *P* = 0.93 without chromosome 4). Similarly, chromosome 4 also entirely accounts for the observed differences in 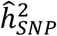 estimates between HE and REML analysis in females (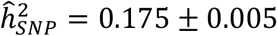 under HE and 0.185 ± 0.005 under REML), as well as the discrepancies between males and females (for males, 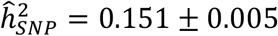 under HE and 0.165 ± 0.006 under REML). Altogether, low polygenicity of urate concentration combined with substantial sex differences in the genetic architecture on chromosome 4 constitute strong departure from assumptions underlying the consistency of HE and REML estimators. There is also prior evidence of female sex hormones effects on urate concentration^38^, which we did not accounted for in our analysis, and further investigations of the sex-specific genetic architecture of the trait are needed. Finally, we detected significant dominance deviation (*P* < 5e-8) at two SNPs on chromosome 1 (top hit for rs12124078, a SNP associated with kidney function^39^) and 4 (larger signal with top hit for rs9998811 located in *SLC2A9* gene region^36^), (**Supplementary Figure 7**) although the total dominance variance explained by these two SNPs remains very small 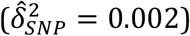.

The lack of evidence for the additive-by-additive variance in our analyses is mostly due to the very large sampling variance of 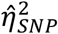. Therefore, potentially real effects on individual traits cannot be ruled out. Nevertheless, we can provide an upper limit for the role of epistatic variance associated with SNPs across the 70 traits of approximately 0.12 (upper bound of the 95% CI from REML analysis, see **Figure 4-A**), based on the SE of the mean 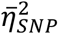 of 0.032. This upper limit (across traits) remains smaller than the well-estimated mean 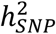 of 0.207. Our power calculations based on REML suggest that ≃2 million unrelated individuals would be necessary to ensure >76% statistical power to detect 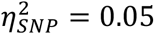 at α = 0.05. The same calculations based on HE regression would only yield a statistical power of 52%. Analysis of such a large sample raise new computational challenges to be addressed in future research.

Besides these limitations, theory and prior evidence also suggest that epistatic variance is likely to be small^7,8,21^, with first order additive-by-additive epistasis expected to be the largest contributor to epistatic variance. In this study, we therefore focused on the additive-by-additive variance and did not estimate higher order interactions or epistatic components involving dominance. Because epistatic GRMs result from the Hadamard products of lower order GRMs, they will quickly tend to identity with increasing order of interactions. It will lead to very large standard errors and estimates of genetic variance indistinguishable from the residual (non-genetic) variance. The same argument holds for any epistatic interaction involving dominance as it will result in even larger standard errors, the variance of the off-diagonal elements of Θ_*D*_ in unrelated European individuals being approximately 10-5, half of the variance of the off-diagonal elements of Θ_*A*_. Finally, Mäki-Tanila and Hill^8^ showed for polygenic traits that when gene-gene interactions (*aa*) are of same magnitude as single-locus effects (*a*), epistatic variance is expected to decrease and eventually disappear with the increase number of causal variants. In this context, epistatic variance lead to mainly additive variance, while dominance is expected to remain the main contributor to non-additive genetic variance. Therefore, and in light of our results showing a lack of dominance variance, that would suggest that epistatic variance is likely to be extremely small in human complex traits.

Our findings can be directly applied to human diseases using a liability threshold model. Yet, it is worth emphasizing that the observed 0-1 scale is expected to show non-additive variance even when variance in liability is fully additive^40^. An analysis of non-additive variance on liability would require the use of generalized non-linear mixed models and would be less powerful than the analysis of quantitative traits. For these reasons we have not attempted to estimate dominance or epistatic variance for liability to common disease in our study. Results for additive SNP-based heritability are by-and-large the same for quantitative traits and common disease, so it seems reasonable to assume that there is likely to be little non-additive variance for liability of disease.

We showed by simulation that we were able to capture a large proportion of the genetic variation from common SNPs in our data even when causal variants were not included. However, because the expected loss of non-additive variance is disproportionally larger than additive variance with the decay of linkage disequilibrium between causal and tagging variants, the contribution of rare variants poorly tagged in our study is expected to be missed. There is evidence that rare variants contribute to narrow sense heritability^41,42^. However, using simulations, Zhu et al.^10^ showed that the observed difference between the additive and dominance variance across traits was unlikely to be explained by a disproportionally missing contribution of rare variants to dominance variation. Moreover, because the amount of non-additive variance also disproportionally depends on allele frequencies as compared to additive variance, with the largest amount expected for intermediate frequencies^8^, contribution of rare variants to the non-additive genetic variance is expected to be minute.

Importantly, the absence of evidence for epistatic variance does not imply the absence of functional epistasis^43,44^. In addition, any significant signal of epistatic variance should be investigated in depth due to potential inferential problems. Indeed, phantom epistasis, that is a non-additive signal generated from incomplete linkage between variants,^45,46^ can induce a bias in the estimates of epistatic variance. A recent study suggested that the effect of phantom epistasis is likely to increase for low density markers which is not necessarily relevant for human data. However, the genetic architecture of a trait, such as local polygenicity (i.e. the non-random spatial distribution of small effect loci) and large effects QTL, could also favor phantom epistasis.

To conclude, the analysis of 70 human complex traits from a large sample of unrelated individuals provides new evidence that genetic variance for complex traits is predominantly additive and suggests negligible dominance variance due to causal variants that are associated with common SNPs. Because of a large standard error, we cannot draw firm conclusions regarding additive-by-additive variance for individual traits, but we can conclude that its upper value is about half of the additive genetic variance captured by common SNPs. We showed that REML lead to substantially larger power as compared to HE at a given sample size, and that sample sizes of many millions of unrelated individuals will be necessary to estimate epistatic variance with sufficient precision.

## Methods

### Genetic model

We assume a model with additive, dominance and epistatic interaction effects (ADAA model). For epistatic variance, we only focus on additive-by-additive interactions. Consider one diploid individual genotyped at L loci, each with major and minor alleles A and B respectively. Let *a*_*i*_ = (*μ*_*AA*_)/2 and *d* = *μ*_*BB*_ − (*μ*_*AA*_ + *μ*_*AA*_)/2, with *μ*_*BB*_, *μ*_*AB*_ and *μ*_*AA*_ being the phenotypic means in the three genotypic classes at one locus. Let *p*_*i*_ be the allele frequency of A at locus *i*, *a*_*i*_ the additive effect, *d*_*i*_ the dominance effect, *aa*_*i,j*_ the additive-by-additive interaction effect between locus *i* and *j*. Under the Hardy-Weinberg equilibrium assumption, we can define the average effect of allele substitution, i.e., 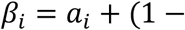 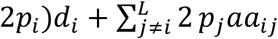, which contains the additive effect, a term due to dominance interaction between two alleles, and a term due to additive-by-additive interactions between pairs of loci^8^. Then, the additive variance at locus 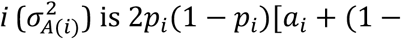 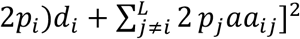, the variance of the average effect of allele substitution, dominance variance 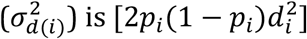, and additive-by-additive variance 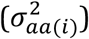 is 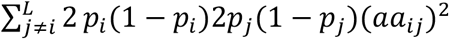. Hence, the genotypic variance 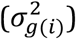 is 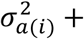 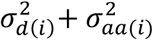.

Let us define the additive genotype coding *x*_*A*((*i*)_ as 0,1 or 2 for genotypes BB, AB and AA respectively, and dominance coding *x*_*D*((*i*)_ as 0, 2*p*_(_ or (4*p*_(_ − 2). This parametrization of *x*_C(()_ following Zhu et al.^10^ ensures the orthogonality with *x*_*A*((*i*)_, compare to the classical 0, 1, 0 code use for the different genotypes, which lead to 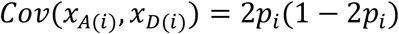. Now including all L biallelic loci, we can fit the additive, dominance and additive-by-additive effect of all SNPs as random effects in a mixed linear model:

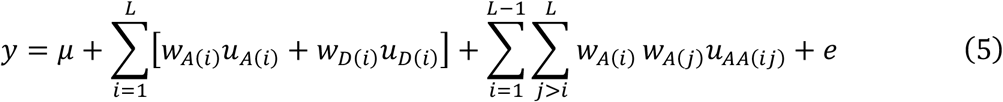

with 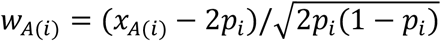 and 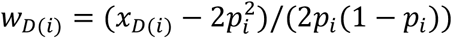 are respectively the standardized form of *x*_*A*_ and *x*_*D*_. *u*_*A*_, *u*_*D*_ and *u*_*AA*_ are the additive, dominance and additive-by-additive random effects of the standardized genotypes, *μ* is the mean term and 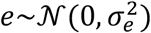 is the residual.

This model can be expressed as an individual based model for a sample of N diploid individuals and the presence of fixed covariates, written in matrix form as:

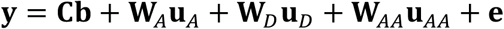

where **y** is a *N* × 1 vector of individuals phenotypes, **W**_*A*_ and **W**_*D*_ are *N* × *L* matrices with one row per individual containing the corresponding *W*_*A*((*i*)_ and *W*_*D*((*i*)_ vectors. **W**_*AA*_ is a 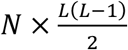 matrix with one row per individual containing all the pairwise *W*_*A*((*i*)_ and *W*_*D*((*j*)_. **u**_**A**_ and **u**_**A**_ are *L* × 1 vectors of locus specific additive and dominance effects, **u**_*AA*_ is a 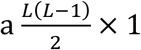 vector of additive-by-additive effects, **e** is an *N* × 1 vector of residuals. **C** is an *N* × *c* matrix of *c* covariates, and **b** is a *c* × 1 vector of corresponding covariate effects.

If we define **g**_*A*_ = **W**_**A**_**u**_**A**_,**g**_**AA**_ = **W**_**AA**_**u**_**AA**_ and **g**_*AA*_ = **W**_*AA*_**u**_*AA*_, then we have:

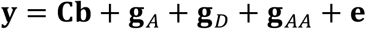

The (co)variance matrix of phenotypes becomes:

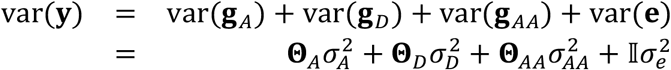

where 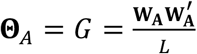 and 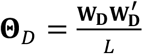 are the additive and dominance genomic relatedness matrices (GRM) as described in Yang et al.^16^ and Zhu et al.^10^, and Θ_***AA***_ is the additive-by-additive genomic relatedness matrix define as:

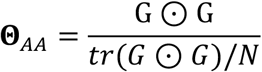

### Genotype data

We analyze a large dataset of 347,849 unrelated (genomic relatedness < 0.05) individuals of European descent (188,088 females and 159,761 males), from the UK Biobank (UKB). Informed consent was obtained from all the subjects and those who expressed the which to be withdrawn have been removed from analysis. We used the release 3 of the UKB where individuals were genotyped on the Affymetrix UK Biobank Axiom array before imputation using the HRC and UK10K reference panel and IMPUTE2^47^. SNPs were filtered for quality control by removing those with missing genotyping rate > 0.05, Hardy-Weinberg equilibrium test *P* < 10^−6^ and minor allele frequency (maf) < 0.01. After filtering, we extracted autosomal HapMap phase 3 (HM3) markers, resulting in 1,130,561 SNPs.

The Additive and Dominance GRM were computed using the standard algorithm of GCTA software^16^ version 1.93.0b as previously described, and the additive-by-additive GRM was computed using R version 3.6.2^48^ from the additive GRM as described in **Equation (2)**. In order to remove any cryptic relatedness, we trimmed the GRMs by removing one individual for each pair with an additive genomic relatedness > 0.025. This resulted in a final dataset of 254,679 unrelated individuals (138,196 females and 116,483 males).

### Phenotypes selection and quality control

We chose 70 continuous trait from the UKB (listed in the **Supplementary Table 3**) with a total number of phenotyped individuals in the all UKB ≥ 60,000, a significant SNP heritability 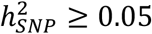 (at *α* = 0.05 level, based on Neale’s lab 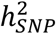 estimates https://nealelab.github.io/UKBB_ldsc/h2_browser.html) and a square pairwise phenotypic correlation *r*^2^ < 0.8. First assessment data only were used, and for phenotypes with left and right measure, only one was chosen randomly. Outlier individuals were removed following Tukey’s method^49^ separately for males and females in every phenotype. This step resulted in a mean and minimum number of phenotyped individuals across traits of 200,550 and 36,690 respectively (among a total of 254,679 individuals). Each sex-specific dataset has been further corrected for age, as well as month of assessment for blood biochemistry traits only, and standardized to z-score. The first 20 eigenvectors of the principal components (PCs), estimated from the 254,679 individuals genotype data using flashPCA2 software^50^, were included as fixed covariates in the REML analyses and the phenotypes were pre-corrected for HE analysis.

### Effective number of phenotypes

For multiple testing correction purpose, we computed the effective number of phenotypes (*P*_*Eff*_) from a general purpose estimator^51^ using the Shannon entropy:

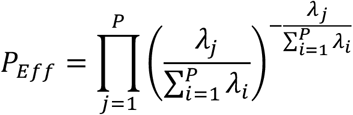

where P is the total number of phenotypes and *λ*_*j*_ the *j*^*th*^ eigenvalue of the phenotypes’ correlation matrix. Using the correlation matrix of our 70 traits, we found *P*_*Eff*_ = 43 and used it for Bonferroni corrections. Because we tested for three variance components but chose only traits with significant SNP-based heritability, we also used two components for the Bonferroni corrections, resulting in a threshold *P-*value of 0.05/(43×2) = 5.8e-4 at *α* = 5%.

### Estimation of genetic variance components

Genetic variance components were estimated using HE regression with phenotypic cross-products and REML. As stated in the main text, HE regression analyses were performed on the full dataset as well as REML when the number of phenotyped individuals was smaller or equal to 60K. Otherwise, we used an inverse-variance weighted (IVW) meta-analysis of 8 sub-datasets of ≃32K individuals. For a single trait, the genetic variance estimate from the IVW meta-analysis of *i* sub-datasets and its associated standard error are:

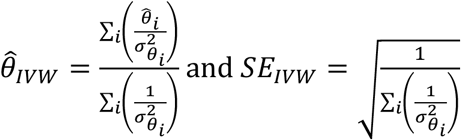

where 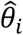 is the genetic variance estimate from the sub-dataset *i*, and 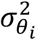 the associated sampling variance. Indications on whether we use a single or meta-analysis for each phenotype can be find in the **Supplementary Table 3**. To obtain unbiased estimates of the variance components, we did not constrain REML estimates to be non-negative. For REML, *P*-values of the estimates were computed as the probability of the likelihood ratio test 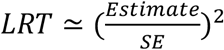 in a Chi-square distribution with one degree of freedom. For HE analysis, we used the reported jackknife SE and *P*-values by GCTA.

We estimated the mean of each variance component estimate 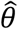, 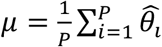 across our P traits, as well as its standard error 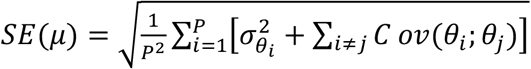 where *Cov*(*θ*_*i*_; *θ*_*j*_) accounts for the non-independence of the traits. We can show from simulations (**Supplementary Figure 8**) that a good approximation of the covariance between genetic estimates is 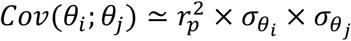 with 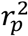 the squared phenotypic correlation between traits *i* and *j*, and 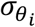 the standard error of the estimate which can be approximate from our analysis (using the reported SE by GCTA). Dominance variance explained by genome-wide significant SNPs was calculated as 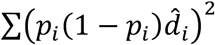 with *p* the empirical allele frequency at SNP *i* and 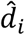 the estimated dominance effect from the genome-wide association analysis. We used R version 3.6.2^48^ to perform the analysis of GCTA outputs.

### Simulation studies

We performed simulation studies to validate our model and theoretical sampling variance of the estimates. We performed simulations based on the real genotypes observed on the 1,130,561 autosomal HM3 SNPs of the 254,679 unrelated European individuals from UKB. Following Zhu et al.^10^, we randomly sampled 100,000 SNPs (across autosomes) as a pool of causal variants and used the remaining SNPs as the observed SNPs. To measure at which extent we were able to capture the additive and dominance variation of causal variants that are associated with common SNPs, we computed the multiple regression *R*2 for the first 10K causal SNPs by regressing the *x*_*A*((*i*)_ (*x*_*D*((*i*)_ under dominance) of the target SNPs with the *x*_*A*((*i*)_ (*x*_*D*((*i*)_ under dominance) of the neighboring SNPs in a 1Mb window.

We generated the phenotypes using a custom C++ program following **Equation (5)** where the additive, dominance, and additive-by-additive effects were generated from a standard normal distribution and adjusted to the expected variance of the additive, dominance and additive-by-additive genome-wide effects (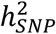, 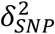 and 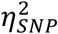). The residuals were then generated from a normal distribution with mean 0 and variance 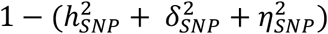. The simulated phenotypes were standardized to z-scores. We chose 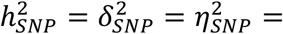 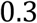 and simulated 100 replicates, each with 1000 randomly sampled causals SNPs from the pool of 100,000 putative causal variants. Missing genotypes were imputed to the mean genotype during the simulation process. The different variance components were estimated using **Equation (1)** under REML and HE regression, including or not causal variants.

To compare results from simulations based on observed genotypes of UKB participants with those under Hardy-Weinberg and linkage equilibrium, we used R v3.6.248 to generate a sample of 35K unrelated individuals genotyped at 100K unlinked markers with uniform allele frequency distributions in the range [0.01;0.99]. Phenotypes were then simulated as described above with the same setting 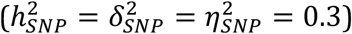. Causal markers were randomly sampled among the 100K unlinked SNPs and all variants were included in the analysis.

## Supporting information

Supplementary Materials for: Estimation of non-additive genetic variance in human complex traits from a large sample of unrelated individuals

## Acknowledgments

We acknowledge funding from the Australian National Health and Medical Research Council (1113400, 1173790) and the Australian Research Council (FT180100186 and FL180100072). This research has been conducted using the UK Biobank Resource under Application Number 12505. We wish to acknowledge The University of Queensland’s Research Computing Centre (RCC) for its support in this research. We also thank Allan McRae for technical support and Jian Zeng for fruitful discussions.

## Author contributions

P.M.V conceived and designed the study. Theory was derived by P.M.V, L.Y and V.H. The UKB phenotypic data were extracted by J.S who also assisted V.H on quality control. V.H and L.Y performed simulations and statistical analysis under the assistance and guidance of P.M.V, N.R.W and J.Y. The manuscript has been written by V.H, L.Y and P.M.V with the participation of all authors. All authors reviewed and approved the final manuscript.

## Web Resources

The URLs for data and software used in this paper are as follow: UK Biobank: https://www.ukbiobank.ac.uk/

FlashPCA2: https://github.com/gabraham/flashpca

GCTA: https://cnsgenomics.com/software/gcta/#Overview

