## Supplementary Materials for: Estimation of non-additive genetic variance in human complex traits from a large sample of unrelated individuals for "Estimation of non-additive genetic variance in human complex traits from a large sample of unrelated individuals"

### Supplementary Figures

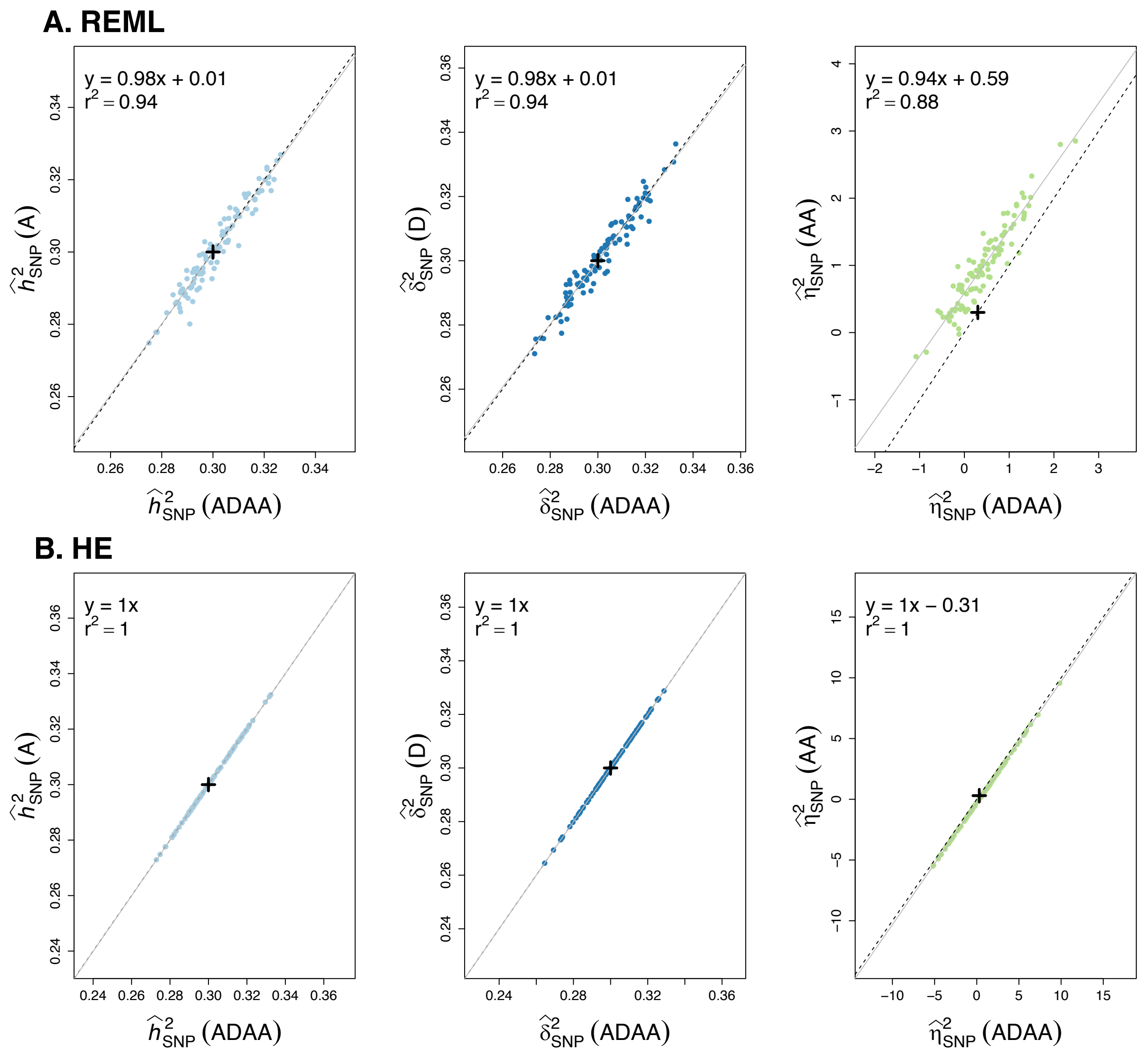


**Supplementary Figure 1: Single-replicate estimates from simulations of unrelated individuals with simulated unlinked markers.** We compared (**A**) REML and (**B**) HE estimates for 100 replicates of simulations of 35,000 unrelated individuals genotyped at 100,000 unlinked markers, when we jointly estimate the different variance components (ADAA) or only one at a time for $h^2_{SNP}$, $\delta^2_{SNP}$ and $\eta^2_{SNP}$. Simulated values are depicted by a black cross, the black dashed line depicts the Y=X line while the solid grey line depicts the linear regression between the corresponding ADAA and single component model.

The squared correlation $r^2$ is close to 1 for all the variance components and $\eta^2_{SNP}$ is the only one showing a large intercept.

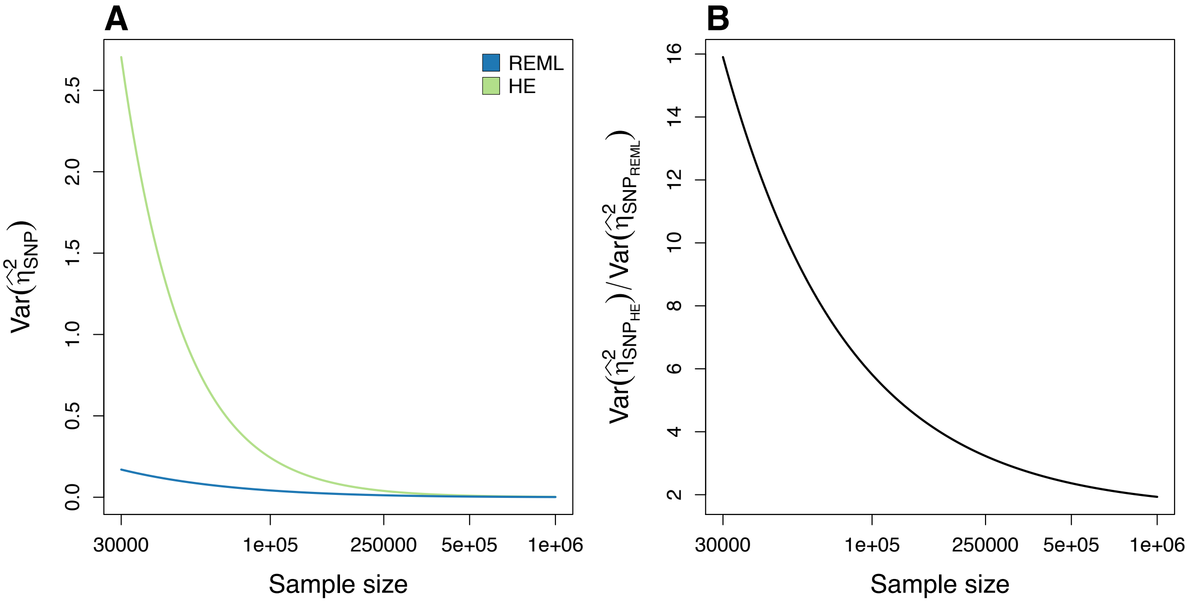


**Supplementary Figure 2: (A) Theoretical $Var(\hat{\eta}^2_{SNP_{REML}})$ (blue) and $Var(\hat{\eta}^2_{SNP_{HE}})$ (green) analysis, and (B) ratio $Var(\hat{\eta}^2_{SNP_{HE}})/Var(\hat{\eta}^2_{SNP_{REML}})$ as a function of sample size in the UKB.** Theoretical expectations have been computed using the observed variance of the elements of the UKB additive GRM of 254,679 unrelated individuals.

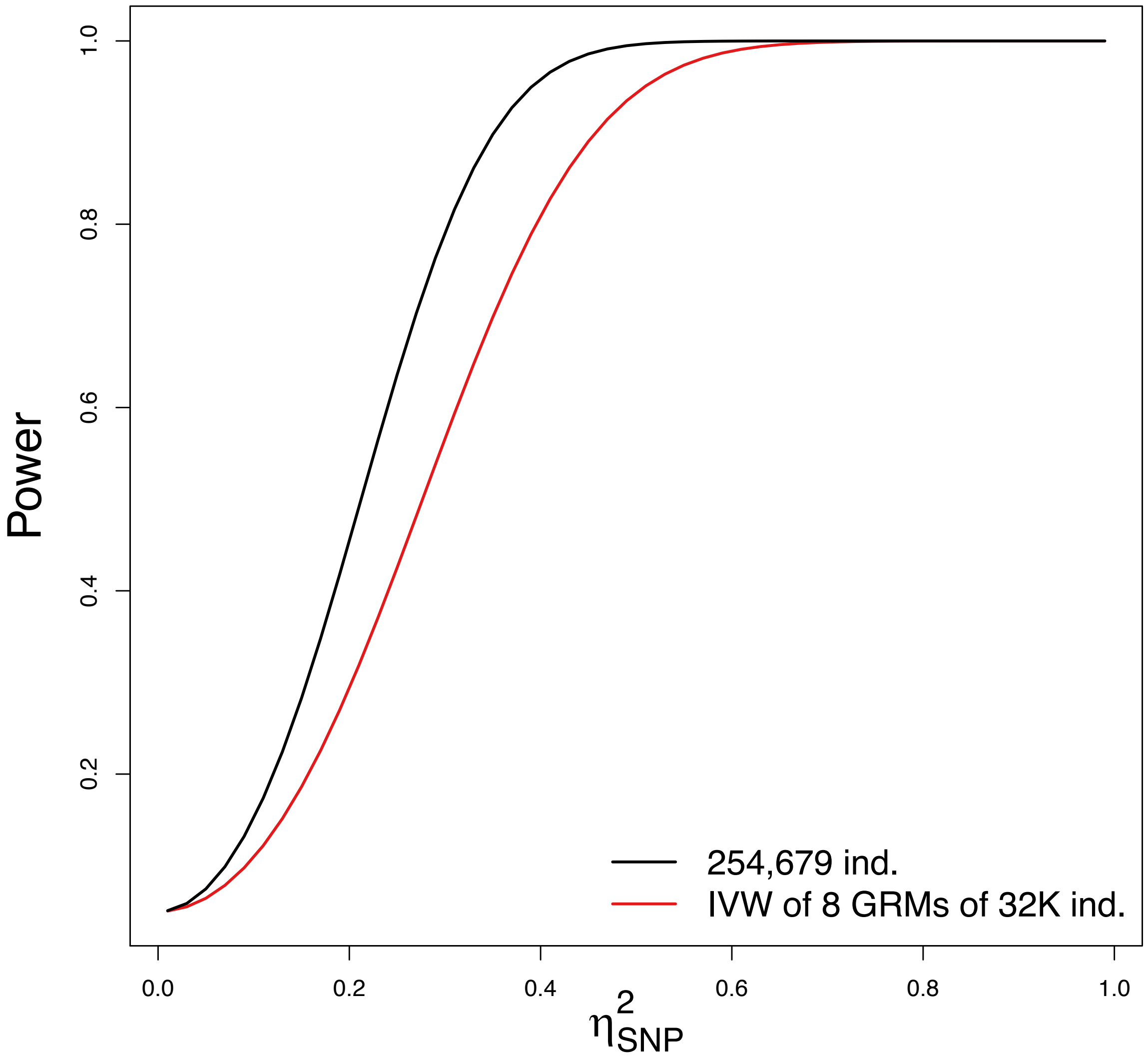


**Supplementary Figure 3: Expected power under REML to detect different amount of $\boldsymbol{\eta^2_{SNP}}$ in the UKB data.** Expected powers are provided for the simultaneous analysis of 254,679 unrelated individuals (solid black line) or for an IVW meta-analysis of 8 sub-datasets of ~32,000 individuals each is performed (solid red line).

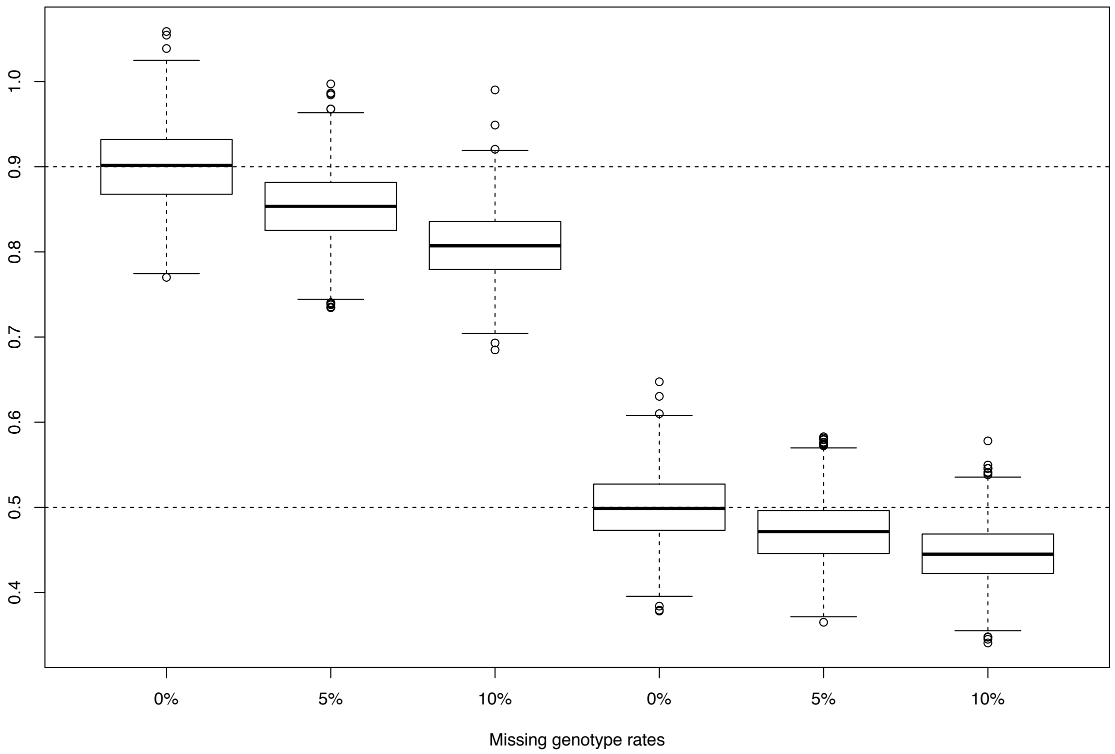


**Supplementary Figure 4 : Effect of genotype missingness on $h^2_{SNP}$ estimates from simulations of unrelated individuals genotyped at unlinked markers.** We simulated 1000 unrelated individuals genotyped at 100 unlinked markers (with population allele frequencies $p$=0.5) with genotype missingness equal to 0%, 5% and 10%. We simulated traits with $h^2_{SNP} = 0.9\ and\ 0.5$ for each scenario using **Equation (5)** of the main manuscript with 1000 replicates each. When simulating phenotypes, the missing genotypes were imputed to the mean genotype. We then computed HE estimates of $h^2_{SNP}$ for each replicate and show that $\mathbb{E}(\hat{h}^2_{SNP}) \simeq (1-r)h^2_{SNP}$, with r the missing genotype rate.

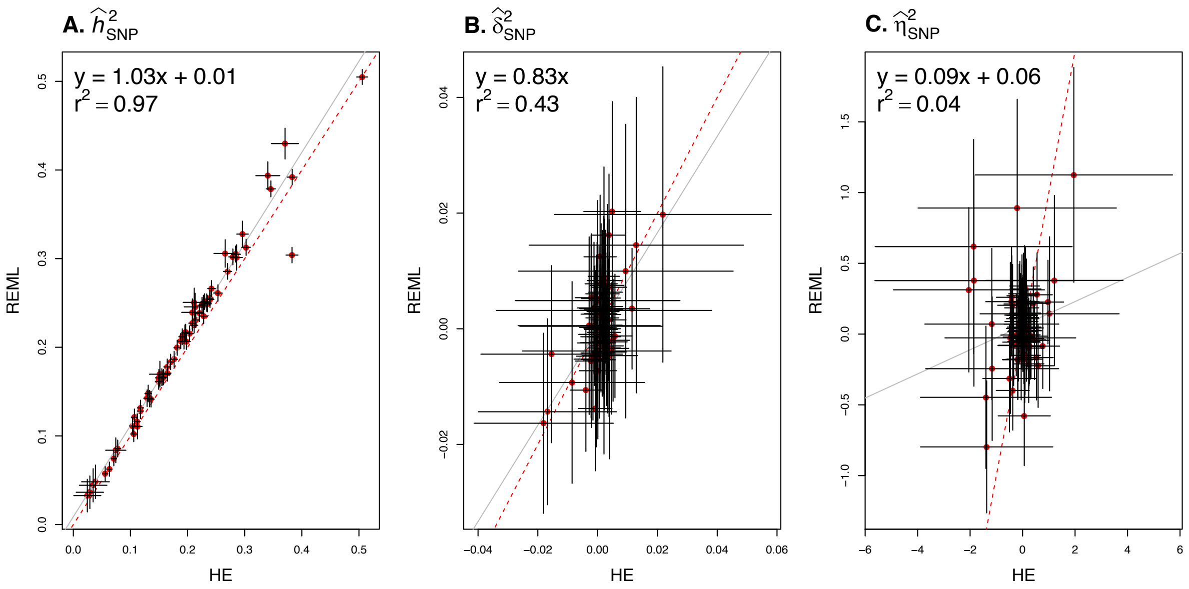


**Supplementary Figure 5: Comparison of variance components estimates from HE regression and GREML for 70 traits measured in UKB participants.** Comparisons between REML and HE are shown for $\hat{h}^2_{SNP}$ (panel A), $\hat{\delta}^2_{SNP}$ (panel B) and $\hat{\eta}^2_{SNP}$ (panel C). For each variance component, we represent the Y=X line (red dashed line) as well as the linear regression between the HE and REML estimates (grey solid line) with its equation and squared correlation ($r^2$). For each estimate, the 95% confidence interval is depicted by a black solid line.

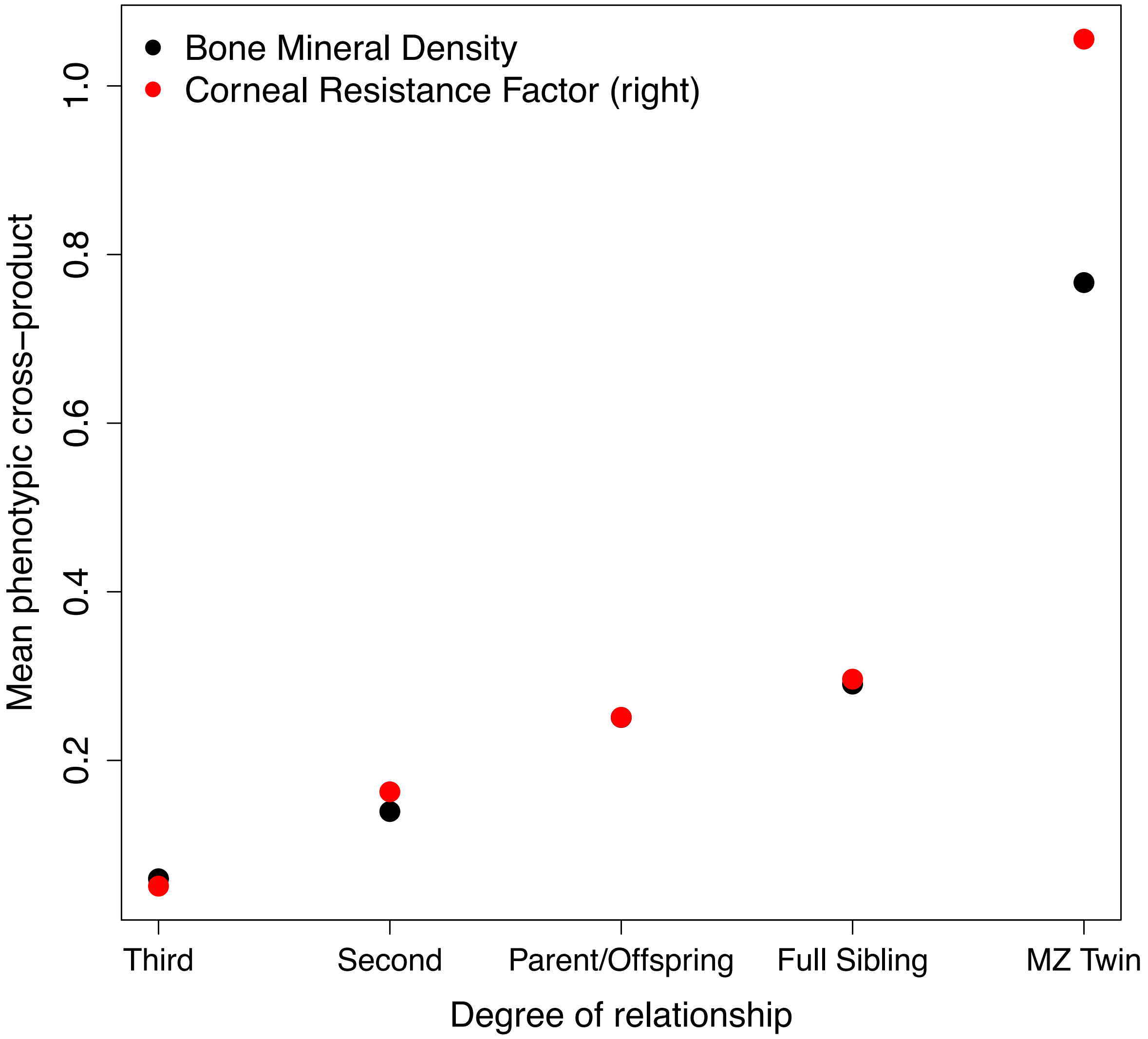


**Supplementary Figure 6: Mean phenotypic cross-product between pairs of individuals for Bone Mineral Density (BMD) and Corneal Resistance Factor (CRF) as a function of family relationship.** We used family data from the UKB and computed the phenotypic cross-products of 44,810 pairs of individuals for BMD (black dots) and 9,375 pairs of individuals for CRF (red dots). In our notations, MZ Twin stands for monozygotic twins.

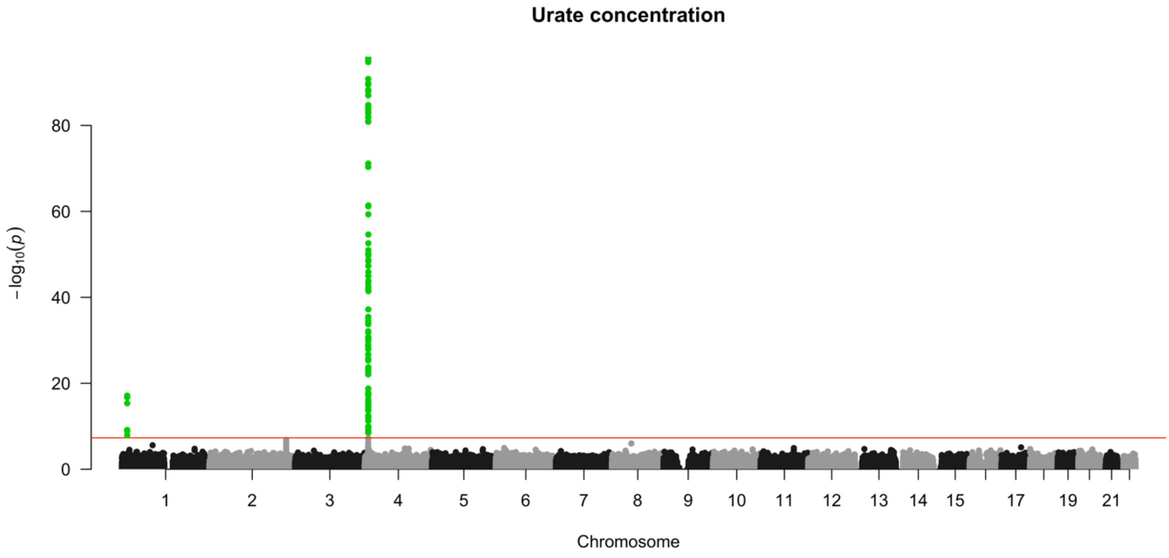


**Supplementary Figure 7: Genome-wide association test for dominance effects for urate concentration.** Manhattan plot showing -log10 of *P*-values for dominance effects from a model fitting both additive and dominance effect with plink2 software. Genome-wide significant SNPs (threshold *P*-value = 5e-8 depicted by the red solid line) are highlighted in green and located within two loci on chromosomes 1 and 4.

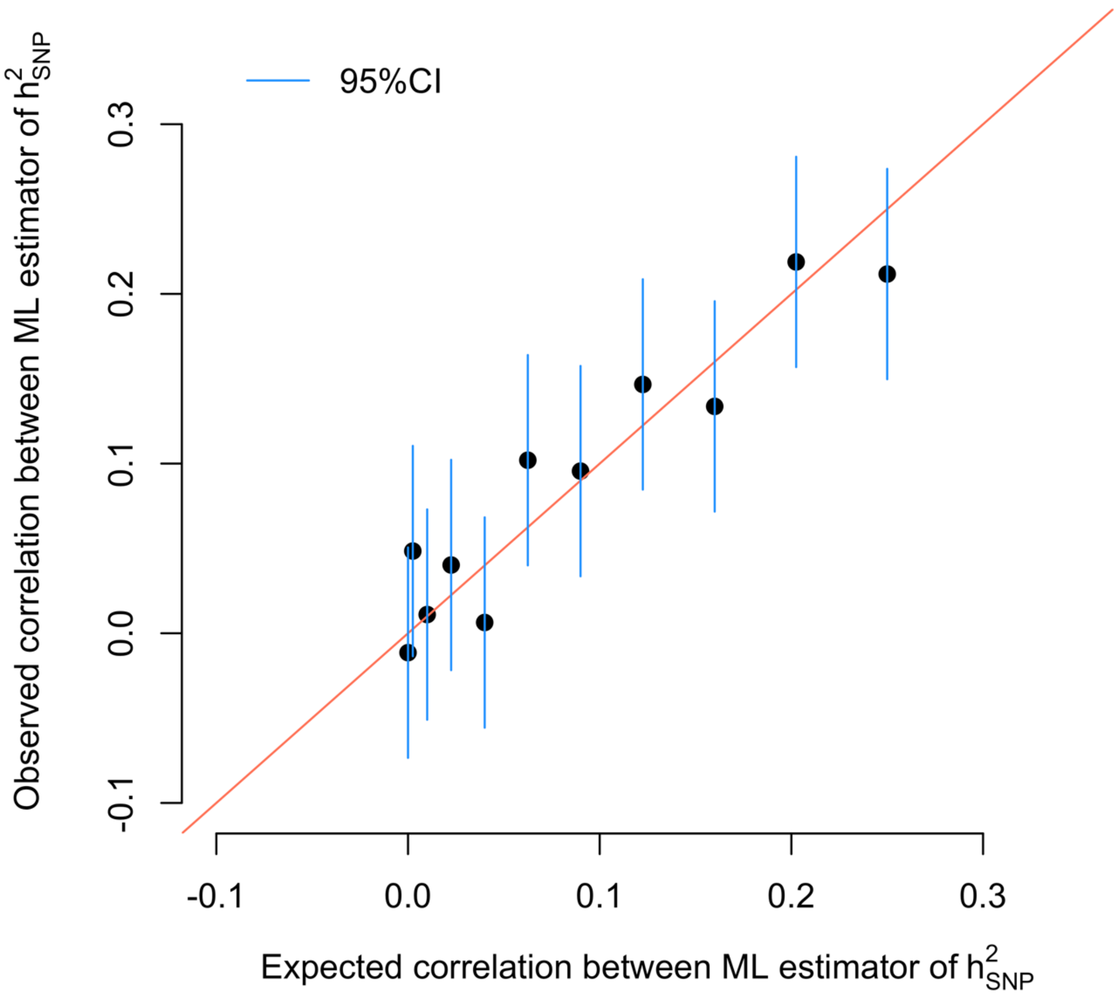


**Supplementary Figure 8 : Observed and expected correlation of Maximum Likelihood (ML) estimates of $h^2_{SNP}$ from simulations of unrelated individuals genotyped at unlinked markers.** We simulated 5000 unrelated individuals genotyped at 5000 unlinked markers (with population allele frequencies *p*=0.5). We simulated pairs of traits with $h^2_{SNP} = 0.5$ using **Equation (5)** of the main manuscript and with phenotypic correlations ranging from 0 to 0.5, with 1000 replicates each. We then computed ML estimates of $h^2_{SNP}$ for each pair of traits and computed, for each simulation setting, the observed correlation between estimates across replicates (blue dots) and compared it to the expected correlation approximated as the squared phenotypic correlation $r_p^2$. The 95% confidence interval of the observed correlations are depicted by the blue solid lines and the Y=X line by the red solid line.

**Supplementary Tables**

**Supplementary Table 1: Mean (variance) of the diagonal and off-diagonal elements of the different GRMs computed from the UKB data.**

| GRM | Off-diagonal | Diagonal |
|---|---|---|
| $\boldsymbol{\Theta}_{A} = \boldsymbol{G}$ | -2.28e-6 (2.03e-5) | 0.99 (8.88e-5) |
| $\boldsymbol{\Theta}_{D}$ | 5.98e-7 (9.31e-6) | 1.00 (2.03e-3) |
| $\boldsymbol{\Theta}_{AA}$ | 2.03e-5 (8.25e-10) | 0.99 (3.65e-4) |

**Supplementary Table 2: Correlation between the off-diagonal (diagonal) elements of the different GRMs computed from the UKB data.**

| GRM | $\boldsymbol{\Theta}_{A} = \boldsymbol{G}$ | $\boldsymbol{\Theta}_{D}$ | $\boldsymbol{\Theta}_{AA}$ |
|---|---|---|---|
| $\boldsymbol{\Theta}_{A} = \boldsymbol{G}$ | 1.00 (1.00) | -2.19e-3 (0.67) | 1.73e-2 (1.00) |
| $\boldsymbol{\Theta}_{D}$ | -2.19e-3 (0.67) | 1.00 (1.00) | 6.24e-3 (0.68) |
| $\boldsymbol{\Theta}_{AA}$ | 1.73e-2 (1.00) | 6.24e-3 (0.68) | 1.00 (1.00) |

**Supplementary Table 3: UKB Phenotypes used for analysis.** For each of the 70 traits, we indicate the UKB code (f.eid), the description of the trait, the remaining number of phenotyped individuals after filtering (N) and whether a single or IVW meta-analysis were performed for REML. We also indicate the different covariates used for each trait, as detailed in the **Methods** section of the main manuscript.

| f.eid | description | N | REML Analysis | Covariates |
|---|---|---|---|---|
| 48 | Waist circumference | 249947 | IVW | Sex, Age, 20 PCs |
| 49 | Hip circumference | 245986 | IVW | Sex, Age, 20 PCs |
| NA | Waist/Hip Ratio | 251349 | IVW | Sex, Age, 20 PCs |
| 50 | Standing height | 251805 | IVW | Sex, Age, 20 PCs |
| 78 | Heel bone mineral density (BMD) T-score, automated | 140801 | IVW | Sex, Age, 20 PCs |
| 3062 | Forced vital capacity (FVC) | 228373 | IVW | Sex, Age, 20 PCs |
| 3143 | Ankle spacing width | 142312 | IVW | Sex, Age, 20 PCs |
| 5084 | Spherical power (right) | 52852 | Single | Sex, Age, 20 PCs |
| 5098 | 6mm weak meridian (right) | 48051 | Single | Sex, Age, 20 PCs |
| 5119 | 3mm cylindrical power (left) | 52752 | Single | Sex, Age, 20 PCs |
| 5135 | 3mm strong meridian (left) | 54535 | Single | Sex, Age, 20 PCs |
| 5254 | Intra-ocular pressure, corneal-compensated (right) | 55467 | Single | Sex, Age, 20 PCs |
| 5255 | Intra-ocular pressure, Goldmann-correlated (right) | 55554 | Single | Sex, Age, 20 PCs |
| 5256 | Corneal hysteresis (right) | 55227 | Single | Sex, Age, 20 PCs |
| 5257 | Corneal resistance factor (right) | 55388 | Single | Sex, Age, 20 PCs |
| 20015 | Sitting height | 250227 | IVW | Sex, Age, 20 PCs |
| 20022 | Birth weight | 138855 | IVW | Sex, Age, 20 PCs |
| 21001 | Body mass index (BMI) | 246468 | IVW | Sex, Age, 20 PCs |
| 23107 | Impedance of leg (right) | 246213 | IVW | Sex, Age, 20 PCs |
| 23110 | Impedance of arm (left) | 247341 | IVW | Sex, Age, 20 PCs |
| 23111 | Leg fat percentage (right) | 247533 | IVW | Sex, Age, 20 PCs |
| 23114 | Leg predicted mass (right) | 244160 | IVW | Sex, Age, 20 PCs |
| 23127 | Trunk fat percentage | 247004 | IVW | Sex, Age, 20 PCs |
| 30010 | Red blood cell (erythrocyte) count | 243594 | IVW | Sex, Age, 20 PCs |
| 30030 | Haematocrit percentage | 243089 | IVW | Sex, Age, 20 PCs |
| 30040 | Mean corpuscular volume | 241538 | IVW | Sex, Age, 20 PCs |
| 30050 | Mean corpuscular haemoglobin | 240123 | IVW | Sex, Age, 20 PCs |
| 30060 | Mean corpuscular haemoglobin concentration | 244229 | IVW | Sex, Age, 20 PCs |
| 30080 | Platelet count | 241800 | IVW | Sex, Age, 20 PCs |
| 30090 | Platelet crit | 241794 | IVW | Sex, Age, 20 PCs |
| 30100 | Mean platelet (thrombocyte) volume | 242375 | IVW | Sex, Age, 20 PCs |
| 30110 | Platelet distribution width | 241762 | IVW | Sex, Age, 20 PCs |
| 30120 | Lymphocyte count | 241089 | IVW | Sex, Age, 20 PCs |
| 30130 | Monocyte count | 237945 | IVW | Sex, Age, 20 PCs |
| 30140 | Neutrophill count | 239906 | IVW | Sex, Age, 20 PCs |
| 30190 | Monocyte percentage | 240599 | IVW | Sex, Age, 20 PCs |
| 30200 | Neutrophill percentage | 243404 | IVW | Sex, Age, 20 PCs |
| 30210 | Eosinophill percentage | 235282 | IVW | Sex, Age, 20 PCs |
| 30250 | Reticulocyte count | 238072 | IVW | Sex, Age, 20 PCs |
| 30260 | Mean reticulocyte volume | 235187 | IVW | Sex, Age, 20 PCs |
| 30270 | Mean sphered cell volume | 239272 | IVW | Sex, Age, 20 PCs |
| 30280 | Immature reticulocyte fraction | 240466 | IVW | Sex, Age, 20 PCs |
| 30290 | High light scatter reticulocyte percentage | 235960 | IVW | Sex, Age, 20 PCs |
| 30510 | Creatinine (enzymatic) in urine | 241281 | IVW | Sex, Age, 20 PCs |
| 30530 | Sodium in urine | 242263 | IVW | Sex, Age, 20 PCs |
| 30600 | Albumin | 219648 | IVW | Sex, Age, month, 20 PCs |
| 30610 | Alkaline phosphatase | 236975 | IVW | Sex, Age, month, 20 PCs |
| 30620 | Alanine aminotransferase | 228870 | IVW | Sex, Age, month, 20 PCs |
| 30640 | Apolipoprotein B | 239036 | IVW | Sex, Age, month, 20 PCs |
| 30650 | Aspartate aminotransferase | 230587 | IVW | Sex, Age, month, 20 PCs |

Follow on next page

**Supplementary Table 3: − (Next)**

| f.eid | description | N | REML Analysis | Covariates |
|---|---|---|---|---|
| 30670 | Urea | 237699 | IVW | Sex, Age, month, 20 PCs |
| 30680 | Calcium | 218275 | IVW | Sex, Age, month, 20 PCs |
| 30690 | Cholesterol | 240572 | IVW | Sex, Age, month, 20 PCs |
| 30710 | C-reactive protein | 220381 | IVW | Sex, Age, month, 20 PCs |
| 30720 | Cystatin C | 235686 | IVW | Sex, Age, month, 20 PCs |
| 30730 | Gamma glutamyltransferase | 221828 | IVW | Sex, Age, month, 20 PCs |
| 30740 | Glucose | 207361 | IVW | Sex, Age, month, 20 PCs |
| 30750 | Glycated haemoglobin (HbA1c) | 230702 | IVW | Sex, Age, month, 20 PCs |
| 30760 | HDL cholesterol | 218172 | IVW | Sex, Age, month, 20 PCs |
| 30770 | IGF-1 | 238147 | IVW | Sex, Age, month, 20 PCs |
| 30810 | Phosphate | 219584 | IVW | Sex, Age, month, 20 PCs |
| 30830 | SHBG | 213743 | IVW | Sex, Age, month, 20 PCs |
| 30860 | Total protein | 218923 | IVW | Sex, Age, month, 20 PCs |
| 30870 | Triglycerides | 232208 | IVW | Sex, Age, month, 20 PCs |
| 30880 | Urate | 238773 | IVW | Sex, Age, month, 20 PCs |
| 30890 | Vitamin D | 230499 | IVW | Sex, Age, month, 20 PCs |
| 100004 | Fat | 36936 | Single | Sex, Age, 20 PCs |
| 100005 | Carbohydrate | 36765 | Single | Sex, Age, 20 PCs |
| 100008 | Total sugars | 36690 | Single | Sex, Age, 20 PCs |
| 100009 | Englyst dietary fibre | 36923 | Single | Sex, Age, 20 PCs |

**Supplementary Table 4: Summary of REML estimates for our 70 UKB traits.** For each trait, we indicate the UKB code (f.eid) and the description. For each estimate of variance component, results are depicted in the format estimate±SE (*P*-value).

| f.eid | description | $\hat{h}^2_{SNP}$ | $\hat{\delta}^2_{SNP}$ | $\hat{\eta}^2_{SNP}$ |
|---|---|---|---|---|
| 48 | Waist circumference | 0.199 ± 4.27e-03 (0.00e+00) | 0.005 ± 5.15e-03 (3.02e-01) | 0.229 ± 1.44e-01 (1.11e-01) |
| 49 | Hip circumference | 0.207 ± 4.34e-03 (0.00e+00) | 0.002 ± 5.21e-03 (6.36e-01) | 0.204 ± 1.44e-01 (1.57e-01) |
| NA | Waist/Hip Ratio | 0.165 ± 4.18e-03 (0.00e+00) | 0.009 ± 5.18e-03 (7.99e-02) | 0.117 ± 1.41e-01 (4.06e-01) |
| 50 | Standing height | 0.505 ± 4.23e-03 (0.00e+00) | 0.004 ± 4.66e-03 (4.10e-01) | 0.217 ± 1.29e-01 (9.09e-02) |
| 78 | Heel bone mineral density (BMD) T-score, automated | 0.328 ± 7.33e-03 (0.00e+00) | 0.003 ± 9.01e-03 (7.00e-01) | -0.578 ± 1.76e-01 (1.03e-03) |
| 3062 | Forced vital capacity (FVC) | 0.261 ± 4.68e-03 (0.00e+00) | 0.001 ± 5.54e-03 (9.11e-01) | 0.109 ± 1.45e-01 (4.53e-01) |
| 3143 | Ankle spacing width | 0.301 ± 7.20e-03 (0.00e+00) | -0.005 ± 8.93e-03 (6.03e-01) | 0.229 ± 1.92e-01 (2.33e-01) |
| 5084 | Spherical power (right) | 0.251 ± 8.01e-03 (8.89e-215) | -0.004 ± 8.29e-03 (6.43e-01) | 0.144 ± 2.72e-01 (5.98e-01) |
| 5098 | 6mm weak meridian (right) | 0.43 ± 8.70e-03 (0.00e+00) | 0.005 ± 8.65e-03 (5.74e-01) | 0.313 ± 2.91e-01 (2.81e-01) |
| 5119 | 3mm cylindrical power (left) | 0.084 ± 7.00e-03 (5.15e-33) | -0.009 ± 8.72e-03 (2.87e-01) | 0.379 ± 3.00e-01 (2.06e-01) |
| 5135 | 3mm strong meridian (left) | 0.394 ± 7.89e-03 (0.00e+00) | -0.004 ± 7.66e-03 (5.67e-01) | 0.071 ± 2.67e-01 (7.91e-01) |
| 5254 | Intra-ocular pressure, corneal-compensated (right) | 0.17 ± 7.30e-03 (1.59e-119) | -0.014 ± 8.04e-03 (7.42e-02) | -0.027 ± 2.67e-01 (9.21e-01) |
| 5255 | Intra-ocular pressure, Goldmann-correlated (right) | 0.239 ± 7.62e-03 (2.05e-216) | -0.016 ± 7.79e-03 (3.60e-02) | -0.447 ± 2.51e-01 (7.55e-02) |
| 5256 | Corneal hysteresis (right) | 0.246 ± 7.68e-03 (2.10e-224) | 0 ± 7.91e-03 (9.56e-01) | -0.244 ± 2.54e-01 (3.36e-01) |
| 5257 | Corneal resistance factor (right) | 0.306 ± 7.77e-03 (0.00e+00) | 0.001 ± 7.68e-03 (9.44e-01) | -0.797 ± 2.33e-01 (6.29e-04) |
| 20015 | Sitting height | 0.392 ± 4.34e-03 (0.00e+00) | 0.005 ± 4.88e-03 (2.86e-01) | -0.082 ± 1.34e-01 (5.39e-01) |
| 20022 | Birth weight | 0.111 ± 7.02e-03 (5.17e-56) | 0.02 ± 9.50e-03 (3.28e-02) | -0.314 ± 1.89e-01 (9.66e-02) |
| 21001 | Body mass index (BMI) | 0.231 ± 4.37e-03 (0.00e+00) | 0.003 ± 5.19e-03 (5.66e-01) | 0.229 ± 1.42e-01 (1.07e-01) |
| 23107 | Impedance of leg (right) | 0.255 ± 4.41e-03 (0.00e+00) | 0.003 ± 5.13e-03 (6.10e-01) | -0.088 ± 1.39e-01 (5.27e-01) |
| 23110 | Impedance of arm (left) | 0.249 ± 4.38e-03 (0.00e+00) | 0.003 ± 5.13e-03 (5.61e-01) | 0.018 ± 1.41e-01 (9.00e-01) |
| 23111 | Leg fat percentage (right) | 0.235 ± 4.34e-03 (0.00e+00) | -0.001 ± 5.13e-03 (7.88e-01) | 0.285 ± 1.43e-01 (4.70e-02) |
| 23114 | Leg predicted mass (right) | 0.286 ± 4.46e-03 (0.00e+00) | 0.001 ± 5.16e-03 (9.22e-01) | 0.213 ± 1.41e-01 (1.30e-01) |
| 23127 | Trunk fat percentage | 0.225 ± 4.34e-03 (0.00e+00) | 0.008 ± 5.21e-03 (1.35e-01) | 0.34 ± 1.42e-01 (1.70e-02) |
| 30010 | Red blood cell (erythrocyte) count | 0.251 ± 4.46e-03 (0.00e+00) | 0.016 ± 5.29e-03 (2.20e-03) | -0.081 ± 1.41e-01 (5.65e-01) |
| 30030 | Haematocrit percentage | 0.183 ± 4.35e-03 (0.00e+00) | 0.012 ± 5.33e-03 (1.96e-02) | -0.079 ± 1.42e-01 (5.79e-01) |
| 30040 | Mean corpuscular volume | 0.302 ± 4.52e-03 (0.00e+00) | 0.002 ± 5.19e-03 (7.17e-01) | -0.21 ± 1.37e-01 (1.25e-01) |
| 30050 | Mean corpuscular haemoglobin | 0.305 ± 4.53e-03 (0.00e+00) | -0.004 ± 5.20e-03 (4.20e-01) | 0.043 ± 1.39e-01 (7.55e-01) |
| 30060 | Mean corpuscular haemoglobin concentration | 0.062 ± 3.99e-03 (3.21e-55) | -0.011 ± 5.31e-03 (4.57e-02) | -0.031 ± 1.42e-01 (8.27e-01) |
| 30080 | Platelet count | 0.305 ± 4.52e-03 (0.00e+00) | -0.003 ± 5.16e-03 (5.66e-01) | -0.04 ± 1.39e-01 (7.72e-01) |
| 30090 | Platelet crit | 0.266 ± 4.49e-03 (0.00e+00) | -0.002 ± 5.21e-03 (6.45e-01) | -0.044 ± 1.39e-01 (7.53e-01) |
| 30100 | Mean platelet (thrombocyte) volume | 0.379 ± 4.54e-03 (0.00e+00) | -0.005 ± 5.05e-03 (2.88e-01) | 0.07 ± 1.37e-01 (6.10e-01) |
| 30110 | Platelet distribution width | 0.247 ± 4.46e-03 (0.00e+00) | -0.005 ± 5.26e-03 (3.15e-01) | 0.271 ± 1.45e-01 (6.18e-02) |
| 30120 | Lymphocyte count | 0.215 ± 4.42e-03 (0.00e+00) | 0.002 ± 5.26e-03 (7.70e-01) | -0.18 ± 1.41e-01 (2.01e-01) |

Follow on next page

**Supplementary Table 4:** – **(Next)**

| f.eid | description | $\hat{h}^2_{SNP}$ | $\hat{\delta}^2_{SNP}$ | $\hat{\eta}^2_{SNP}$ |
|---|---|---|---|---|
| 30130 | Monocyte count | 0.216 ± 4.51e-03 (0.00e+00) | 0 ± 5.39e-03 (9.67e-01) | -0.006 ± 1.44e-01 (9.64e-01) |
| 30140 | Neutrophill count | 0.178 ± 4.38e-03 (0.00e+00) | -0.001 ± 5.33e-03 (9.07e-01) | 0.278 ± 1.48e-01 (6.07e-02) |
| 30190 | Monocyte percentage | 0.212 ± 4.46e-03 (0.00e+00) | -0.005 ± 5.28e-03 (3.16e-01) | -0.181 ± 1.41e-01 (1.99e-01) |
| 30200 | Neutrophill percentage | 0.167 ± 4.29e-03 (0.00e+00) | -0.002 ± 5.28e-03 (7.06e-01) | 0.024 ± 1.42e-01 (8.67e-01) |
| 30210 | Eosinophill percentage | 0.186 ± 4.48e-03 (0.00e+00) | 0.005 ± 5.52e-03 (3.30e-01) | -0.178 ± 1.44e-01 (2.14e-01) |
| 30250 | Reticulocyte count | 0.206 ± 4.46e-03 (0.00e+00) | 0.005 ± 5.41e-03 (3.67e-01) | 0.142 ± 1.46e-01 (3.31e-01) |
| 30260 | Mean reticulocyte volume | 0.217 ± 4.55e-03 (0.00e+00) | 0.006 ± 5.47e-03 (2.69e-01) | 0.319 ± 1.46e-01 (2.90e-02) |
| 30270 | Mean sphered cell volume | 0.227 ± 4.49e-03 (0.00e+00) | 0.006 ± 5.35e-03 (2.37e-01) | 0.132 ± 1.43e-01 (3.55e-01) |
| 30280 | Immature reticulocyte fraction | 0.161 ± 4.33e-03 (6.29e-303) | -0.003 ± 5.37e-03 (5.37e-01) | -0.099 ± 1.44e-01 (4.90e-01) |
| 30290 | High light scatter reticulocyte percentage | 0.207 ± 4.49e-03 (0.00e+00) | -0.001 ± 5.41e-03 (8.00e-01) | -0.044 ± 1.45e-01 (7.60e-01) |
| 30510 | Creatinine (enzymatic) in urine | 0.057 ± 4.03e-03 (6.55e-46) | 0 ± 5.41e-03 (9.74e-01) | -0.021 ± 1.45e-01 (8.83e-01) |
| 30530 | Sodium in urine | 0.074 ± 4.06e-03 (6.72e-75) | -0.002 ± 5.37e-03 (6.50e-01) | 0.226 ± 1.48e-01 (1.26e-01) |
| 30600 | Albumin | 0.148 ± 4.69e-03 (5.44e-218) | 0.003 ± 5.88e-03 (6.67e-01) | -0.015 ± 1.49e-01 (9.22e-01) |
| 30610 | Alkaline phosphatase | 0.249 ± 4.55e-03 (0.00e+00) | -0.001 ± 5.35e-03 (8.17e-01) | 0.148 ± 1.42e-01 (2.96e-01) |
| 30620 | Alanine aminotransferase | 0.102 ± 4.38e-03 (3.10e-120) | -0.003 ± 5.68e-03 (5.50e-01) | -0.052 ± 1.50e-01 (7.29e-01) |
| 30640 | Apolipoprotein B | 0.132 ± 4.32e-03 (1.98e-204) | 0 ± 5.42e-03 (9.83e-01) | 0.04 ± 1.44e-01 (7.80e-01) |
| 30650 | Aspartate aminotransferase | 0.141 ± 4.45e-03 (2.40e-221) | -0.004 ± 5.60e-03 (4.73e-01) | 0.134 ± 1.47e-01 (3.61e-01) |
| 30670 | Urea | 0.116 ± 4.28e-03 (1.74e-162) | -0.014 ± 5.39e-03 (1.05e-02) | 0.26 ± 1.47e-01 (7.65e-02) |
| 30680 | Calcium | 0.142 ± 4.70e-03 (2.72e-202) | -0.005 ± 5.92e-03 (3.75e-01) | 0.049 ± 1.53e-01 (7.47e-01) |
| 30690 | Cholesterol | 0.128 ± 4.25e-03 (3.43e-199) | -0.006 ± 5.34e-03 (2.96e-01) | 0.228 ± 1.45e-01 (1.15e-01) |
| 30710 | C-reactive protein | 0.142 ± 4.64e-03 (5.22e-206) | 0 ± 5.85e-03 (9.98e-01) | 0.089 ± 1.52e-01 (5.61e-01) |
| 30720 | Cystatin C | 0.313 ± 4.58e-03 (0.00e+00) | 0.008 ± 5.31e-03 (1.17e-01) | -0.057 ± 1.38e-01 (6.80e-01) |
| 30730 | Gamma glutamyltransferase | 0.166 ± 4.69e-03 (5.00e-273) | 0.007 ± 5.86e-03 (2.21e-01) | -0.398 ± 1.40e-01 (4.49e-03) |
| 30740 | Glucose | 0.086 ± 4.76e-03 (1.71e-72) | 0.003 ± 6.32e-03 (6.02e-01) | -0.177 ± 1.53e-01 (2.48e-01) |
| 30750 | Glycated haemoglobin (HbA1c) | 0.238 ± 4.64e-03 (0.00e+00) | -0.001 ± 5.52e-03 (8.61e-01) | 0.027 ± 1.45e-01 (8.55e-01) |
| 30760 | HDL cholesterol | 0.249 ± 4.90e-03 (0.00e+00) | -0.002 ± 5.79e-03 (6.72e-01) | 0.225 ± 1.53e-01 (1.41e-01) |
| 30770 | IGF-1 | 0.254 ± 4.51e-03 (0.00e+00) | 0.008 ± 5.35e-03 (1.60e-01) | -0.057 ± 1.42e-01 (6.86e-01) |
| 30810 | Phosphate | 0.121 ± 4.62e-03 (3.30e-151) | 0.001 ± 5.92e-03 (9.24e-01) | -0.166 ± 1.50e-01 (2.67e-01) |
| 30830 | SHBG | 0.25 ± 4.98e-03 (0.00e+00) | -0.007 ± 5.90e-03 (2.19e-01) | 0.185 ± 1.52e-01 (2.22e-01) |
| 30860 | Total protein | 0.166 ± 4.74e-03 (1.04e-269) | -0.002 ± 5.86e-03 (6.99e-01) | -0.085 ± 1.50e-01 (5.71e-01) |
| 30870 | Triglycerides | 0.171 ± 4.49e-03 (1.83e-316) | -0.005 ± 5.54e-03 (3.92e-01) | -0.005 ± 1.44e-01 (9.71e-01) |
| 30880 | Urate | 0.304 ± 4.37e-03 (0.00e+00) | 0.003 ± 5.22e-03 (5.07e-01) | -0.15 ± 1.39e-01 (2.82e-01) |
| 30890 | Vitamin D | 0.111 ± 4.36e-03 (1.37e-143) | 0.004 ± 5.68e-03 (5.25e-01) | -0.224 ± 1.47e-01 (1.29e-01) |
| 100004 | Fat | 0.036 ± 9.22e-03 (7.88e-05) | 0.003 ± 1.24e-02 (7.98e-01) | 1.125 ± 3.80e-01 (3.03e-03) |
| 100005 | Carbohydrate | 0.044 ± 9.38e-03 (2.32e-06) | 0.01 ± 1.27e-02 (4.33e-01) | 0.891 ± 3.84e-01 (2.04e-02) |

Follow on next page

**Supplementary Table 4: – (Next)**

| f.eid | description | $\hat{h}^2_{SNP}$ | $\hat{\delta}^2_{SNP}$ | $\hat{\eta}^2_{SNP}$ |
|---|---|---|---|---|
| 100008 | Total sugars | 0.048 ± 9.43e-03 (3.10e-07) | 0.02 ± 1.28e-02 (1.22e-01) | 0.378 ± 3.73e-01 (3.11e-01) |
| 100009 | Englyst dietary fibre | 0.033 ± 9.19e-03 (3.79e-04) | 0.014 ± 1.28e-02 (2.58e-01) | 0.618 ± 3.79e-01 (1.03e-01) |

**Supplementary Table 5: Summary of HE estimates for our 70 UKB traits.** For each trait, we indicate the UKB code (f.eid) and the description. For each estimate of variance component, results are depicted in the format estimate±SE (*P*-value).

| f.eid | description | $\hat{h}^2_{SNP}$ | $\hat{\delta}^2_{SNP}$ | $\hat{\eta}^2_{SNP}$ |
|---|---|---|---|---|
| 48 | Waist circumference | 0.182±2.98e-03 (0.00e+00) | 0.002±2.66e-03 (3.90e-01) | 0.066±2.80e-01 (8.14e-01) |
| 49 | Hip circumference | 0.188±3.07e-03 (0.00e+00) | 0.002±2.66e-03 (3.57e-01) | -0.382±2.83e-01 (1.77e-01) |
| NA | Waist/Hip Ratio | 0.157±2.81e-03 (0.00e+00) | 0.002±2.63e-03 (4.28e-01) | 0.051±2.78e-01 (8.55e-01) |
| 50 | Standing height | 0.506±4.84e-03 (0.00e+00) | 0.003±2.63e-03 (2.27e-01) | 0.408±2.87e-01 (1.54e-01) |
| 78 | Heel bone mineral density (BMD) T-score, automated | 0.296±5.08e-03 (0.00e+00) | 0.003±4.63e-03 (4.77e-01) | 0.063±4.97e-01 (8.98e-01) |
| 3062 | Forced vital capacity (FVC) | 0.253±3.66e-03 (0.00e+00) | 0.002±2.91e-03 (4.09e-01) | 0.005±3.09e-01 (9.88e-01) |
| 3143 | Ankle spacing width | 0.286±5.10e-03 (0.00e+00) | 0.004±4.61e-03 (3.81e-01) | -0.422±4.91e-01 (3.90e-01) |
| 5084 | Spherical power (right) | 0.212±9.70e-03 (1.57e-105) | 0±1.25e-02 (9.79e-01) | 1.028±1.32e+00 (4.38e-01) |
| 5098 | 6mm weak meridian (right) | 0.371±1.19e-02 (2.37e-213) | 0±1.38e-02 (1.00e+00) | -2.047±1.44e+00 (1.55e-01) |
| 5119 | 3mm cylindrical power (left) | 0.074±8.87e-03 (5.45e-17) | -0.009±1.21e-02 (4.82e-01) | 1.212±1.31e+00 (3.55e-01) |
| 5135 | 3mm strong meridian (left) | 0.34±1.05e-02 (3.50e-229) | -0.015±1.18e-02 (1.94e-01) | -1.172±1.28e+00 (3.58e-01) |
| 5254 | Intra-ocular pressure, corneal-compensated (right) | 0.152±9.06e-03 (9.00e-63) | -0.017±1.16e-02 (1.47e-01) | -0.466±1.25e+00 (7.08e-01) |
| 5255 | Intra-ocular pressure, Goldmann-correlated (right) | 0.208±9.40e-03 (5.54e-109) | -0.018±1.16e-02 (1.21e-01) | -1.395±1.25e+00 (2.64e-01) |
| 5256 | Corneal hysteresis (right) | 0.213±9.55e-03 (1.64e-110) | -0.002±1.19e-02 (8.55e-01) | -1.159±1.27e+00 (3.60e-01) |
| 5257 | Corneal resistance factor (right) | 0.266±9.91e-03 (1.59e-158) | -0.003±1.19e-02 (8.11e-01) | -1.368±1.26e+00 (2.78e-01) |
| 20015 | Sitting height | 0.383±4.28e-03 (0.00e+00) | 0±2.62e-03 (9.46e-01) | 0.255±2.87e-01 (3.75e-01) |
| 20022 | Birth weight | 0.112±4.01e-03 (3.16e-173) | 0.005±4.76e-03 (3.04e-01) | -0.501±5.06e-01 (3.22e-01) |
| 21001 | Body mass index (BMI) | 0.214±3.22e-03 (0.00e+00) | 0.002±2.67e-03 (5.09e-01) | -0.103±2.85e-01 (7.19e-01) |
| 23107 | Impedance of leg (right) | 0.239±3.36e-03 (0.00e+00) | 0.001±2.67e-03 (7.35e-01) | 0.248±2.89e-01 (3.90e-01) |
| 23110 | Impedance of arm (left) | 0.234±3.38e-03 (0.00e+00) | 0.002±2.65e-03 (4.78e-01) | 0.008±2.87e-01 (9.77e-01) |
| 23111 | Leg fat percentage (right) | 0.228±3.35e-03 (0.00e+00) | 0±2.67e-03 (9.11e-01) | 0.036±2.84e-01 (9.00e-01) |
| 23114 | Leg predicted mass (right) | 0.27±3.59e-03 (0.00e+00) | -0.001±2.65e-03 (6.70e-01) | -0.131±2.89e-01 (6.52e-01) |
| 23127 | Trunk fat percentage | 0.211±3.20e-03 (0.00e+00) | 0.001±2.69e-03 (6.65e-01) | 0.077±2.83e-01 (7.85e-01) |
| 30010 | Red blood cell (erythrocyte) count | 0.23±3.37e-03 (0.00e+00) | 0.004±2.73e-03 (1.64e-01) | -0.366±2.88e-01 (2.04e-01) |
| 30030 | Haematocrit percentage | 0.17±3.05e-03 (0.00e+00) | 0.001±2.71e-03 (7.59e-01) | -0.342±2.88e-01 (2.35e-01) |
| 30040 | Mean corpuscular volume | 0.279±3.65e-03 (0.00e+00) | 0.002±2.76e-03 (5.66e-01) | 0.233±2.91e-01 (4.25e-01) |
| 30050 | Mean corpuscular haemoglobin | 0.285±3.72e-03 (0.00e+00) | 0±2.76e-03 (9.90e-01) | 0.208±2.93e-01 (4.78e-01) |
| 30060 | Mean corpuscular haemoglobin concentration | 0.063±2.28e-03 (2.28e-168) | -0.004±2.64e-03 (1.37e-01) | 0.235±2.85e-01 (4.11e-01) |
| 30080 | Platelet count | 0.284±3.63e-03 (0.00e+00) | 0.003±2.78e-03 (2.75e-01) | 0.295±2.90e-01 (3.09e-01) |
| 30090 | Platelet crit | 0.242±3.37e-03 (0.00e+00) | 0±2.74e-03 (9.39e-01) | 0.452±2.90e-01 (1.20e-01) |
| 30100 | Mean platelet (thrombocyte) volume | 0.346±3.91e-03 (0.00e+00) | 0.001±2.69e-03 (8.34e-01) | -0.127±2.89e-01 (6.59e-01) |
| 30110 | Platelet distribution width | 0.225±3.31e-03 (0.00e+00) | 0±2.70e-03 (9.85e-01) | -0.021±2.87e-01 (9.41e-01) |
| 30120 | Lymphocyte count | 0.204±3.29e-03 (0.00e+00) | 0.004±2.94e-03 (2.19e-01) | -0.176±2.92e-01 (5.46e-01) |

Follow on next page

**Supplementary Table 5: – (Next)**

| f.eid | description | $\hat{h}^2_{SNP}$ | $\hat{\delta}^2_{SNP}$ | $\hat{\eta}^2_{SNP}$ |
|---|---|---|---|---|
| 30130 | Monocyte count | 0.195±3.13e-03 (0.00e+00) | 0±2.76e-03 (8.59e-01) | -0.24±2.93e-01 (4.12e-01) |
| 30140 | Neutrophill count | 0.164±3.02e-03 (0.00e+00) | 0.002±2.77e-03 (4.56e-01) | 0.552±2.91e-01 (5.80e-02) |
| 30190 | Monocyte percentage | 0.191±3.07e-03 (0.00e+00) | 0±2.73e-03 (9.91e-01) | -0.188±2.89e-01 (5.15e-01) |
| 30200 | Neutrophill percentage | 0.158±2.96e-03 (0.00e+00) | 0.005±2.80e-03 (6.53e-02) | 0.156±2.87e-01 (5.88e-01) |
| 30210 | Eosinophill percentage | 0.176±3.13e-03 (0.00e+00) | -0.002±2.73e-03 (4.48e-01) | -0.054±2.96e-01 (8.55e-01) |
| 30250 | Reticulocyte count | 0.198±3.34e-03 (0.00e+00) | 0.003±2.75e-03 (3.29e-01) | 0.091±2.93e-01 (7.55e-01) |
| 30260 | Mean reticulocyte volume | 0.197±3.17e-03 (0.00e+00) | 0.002±2.78e-03 (5.63e-01) | 0.11±2.96e-01 (7.11e-01) |
| 30270 | Mean sphered cell volume | 0.209±3.24e-03 (0.00e+00) | 0.003±2.81e-03 (3.47e-01) | -0.014±2.91e-01 (9.62e-01) |
| 30280 | Immature reticulocyte fraction | 0.149±2.96e-03 (0.00e+00) | 0.002±2.72e-03 (4.21e-01) | 0.238±2.91e-01 (4.14e-01) |
| 30290 | High light scatter reticulocyte percentage | 0.194±3.31e-03 (0.00e+00) | 0.002±2.77e-03 (4.33e-01) | 0.152±2.95e-01 (6.06e-01) |
| 30510 | Creatinine (enzymatic) in urine | 0.056±2.25e-03 (9.23e-136) | 0.001±2.73e-03 (7.70e-01) | -0.162±2.90e-01 (5.77e-01) |
| 30530 | Sodium in urine | 0.071±2.40e-03 (3.80e-192) | 0.001±2.74e-03 (8.40e-01) | 0.968±2.97e-01 (1.13e-03) |
| 30600 | Albumin | 0.131±2.93e-03 (0.00e+00) | 0±3.01e-03 (8.73e-01) | 0.322±3.19e-01 (3.13e-01) |
| 30610 | Alkaline phosphatase | 0.235±3.40e-03 (0.00e+00) | 0.006±2.85e-03 (4.18e-02) | 0.099±2.96e-01 (7.38e-01) |
| 30620 | Alanine aminotransferase | 0.106±2.71e-03 (0.00e+00) | 0.005±2.90e-03 (1.08e-01) | 0.05±3.05e-01 (8.69e-01) |
| 30640 | Apolipoprotein B | 0.118±2.62e-03 (0.00e+00) | 0.001±2.76e-03 (6.85e-01) | -0.036±2.92e-01 (9.02e-01) |
| 30650 | Aspartate aminotransferase | 0.136±2.88e-03 (0.00e+00) | 0.004±2.87e-03 (1.84e-01) | -0.075±3.02e-01 (8.04e-01) |
| 30670 | Urea | 0.112±2.67e-03 (0.00e+00) | -0.001±2.80e-03 (7.79e-01) | -0.41±2.94e-01 (1.63e-01) |
| 30680 | Calcium | 0.129±2.94e-03 (0.00e+00) | -0.002±2.96e-03 (5.03e-01) | 0.059±3.22e-01 (8.54e-01) |
| 30690 | Cholesterol | 0.118±2.71e-03 (0.00e+00) | -0.002±2.70e-03 (4.28e-01) | 0.063±2.90e-01 (8.28e-01) |
| 30710 | C-reactive protein | 0.134±2.90e-03 (0.00e+00) | 0.002±3.04e-03 (6.15e-01) | -0.041±3.16e-01 (8.97e-01) |
| 30720 | Cystatin C | 0.302±3.83e-03 (0.00e+00) | 0.002±2.86e-03 (4.34e-01) | -0.161±2.97e-01 (5.87e-01) |
| 30730 | Gamma glutamyltransferase | 0.15±2.97e-03 (0.00e+00) | 0.004±2.96e-03 (1.95e-01) | -0.374±3.16e-01 (2.35e-01) |
| 30740 | Glucose | 0.078±2.67e-03 (2.38e-186) | 0.002±3.21e-03 (5.36e-01) | 0.083±3.36e-01 (8.06e-01) |
| 30750 | Glycated haemoglobin (HbA1c) | 0.221±3.44e-03 (0.00e+00) | -0.001±2.83e-03 (8.15e-01) | 0.077±3.04e-01 (8.01e-01) |
| 30760 | HDL cholesterol | 0.228±3.44e-03 (0.00e+00) | -0.001±3.04e-03 (8.41e-01) | 0.145±3.20e-01 (6.51e-01) |
| 30770 | IGF-1 | 0.239±3.47e-03 (0.00e+00) | 0.003±2.78e-03 (2.48e-01) | 0.086±2.93e-01 (7.69e-01) |
| 30810 | Phosphate | 0.107±2.75e-03 (0.00e+00) | 0.002±2.97e-03 (4.40e-01) | 0.532±3.19e-01 (9.57e-02) |
| 30830 | SHBG | 0.234±3.63e-03 (0.00e+00) | 0±3.08e-03 (9.79e-01) | -0.031±3.25e-01 (9.23e-01) |
| 30860 | Total protein | 0.155±3.11e-03 (0.00e+00) | 0.004±3.02e-03 (2.01e-01) | 0.769±3.20e-01 (1.63e-02) |
| 30870 | Triglycerides | 0.165±3.00e-03 (0.00e+00) | -0.002±2.80e-03 (5.77e-01) | 0.069±3.01e-01 (8.18e-01) |
| 30880 | Urate | 0.383±5.27e-03 (0.00e+00) | 0.012±2.95e-03 (9.58e-05) | 0.094±2.93e-01 (7.49e-01) |
| 30890 | Vitamin D | 0.104±2.71e-03 (1.98e-321) | -0.001±2.81e-03 (6.65e-01) | 0.592±3.07e-01 (5.40e-02) |
| 100004 | Fat | 0.029±1.20e-02 (1.56e-02) | 0.002±1.80e-02 (9.06e-01) | 1.953±1.88e+00 (2.99e-01) |
| 100005 | Carbohydrate | 0.035±1.20e-02 (3.99e-03) | 0.009±1.80e-02 (6.00e-01) | -0.205±1.89e+00 (9.13e-01) |

Follow on next page

**Supplementary Table 5: – (Next)**

| f.eid | description | $\hat{h}^2_{SNP}$ | $\hat{\delta}^2_{SNP}$ | $\hat{\eta}^2_{SNP}$ |
|---|---|---|---|---|
| 100008 | Total sugars | 0.039±1.22e-02 (1.41e-03) | 0.022±1.81e-02 (2.28e-01) | -1.854±1.89e+00 (3.26e-01) |
| 100009 | Englyst dietary fibre | 0.025±1.19e-02 (3.89e-02) | 0.013±1.79e-02 (4.71e-01) | -1.866±1.88e+00 (3.20e-01) |

### Supplementary Note 1

#### Effect of sampled allele frequencies on the lack of orthogonality of the $\eta^2_{SNP}$ estimates

Using simulated data based on observed genotypes of 254,679 UKB participants, the model described in the Equation 1 of the main manuscript showed good orthogonal properties for $h^2_{SNP}$ and $\delta^2_{SNP}$ estimates (fitting all the GRMs or only one at a time did not change the estimates, **Figure 1**). However, a deviation was observed between estimates of $\eta^2_{SNP}$ in the ADAA model (fitting jointly the additive, dominance and additive-by-additive GRMs) or when $\Theta_{AA}$ only was fit (resulting in biased estimates of $\eta^2_{SNP}$, t-test *P*-value<2.2e-16). This is expected when LD is present in the data, as orthogonal epistatic estimates cannot be obtained. However, we observed the same phenomenon using simulations of unrelated individuals genotyped at unlinked markers (**Supplementary Figure 1**). This problem is hard to investigate under REML, however, it is more computationally tractable under HE regression framework and we will demonstrate that it can be explained because of a sampling effect where we used estimated allele frequencies for a particular dataset instead of the true population frequencies when computing the different GRM.

Assume the same model as described in the **Methods** section of the main manuscript where a trait $y$ is genetically partitioned into additive, dominance and additive-by-additive effects, the HE regression model using the phenotypic cross-products is:

$$y_i y_j = b_0 + \Theta_{A_{ij}}\sigma^2_A + \Theta_{D_{ij}}\sigma^2_D + \Theta_{AA_{ij}}\sigma^2_{AA} + e_{ij}$$

with $y_i$ the phenotypic value for individual *i* while $\Theta_{A_{ij}}$, $\Theta_{D_{ij}}$ and $\Theta_{AA_{ij}}$ are the off-diagonal elements of the additive, dominance and additive-by-additive GRM respectively. In a multiple regression where the three GRMs are fitted, partial regression coefficients are estimated and the covariances between regressors are controlled such that confounding effects are avoided, leading to $\mathbb{E}(\hat{\theta}) = \theta$, with $\theta$ the regressor. However, when fitting a simple linear regression, marginal regression coefficients are estimated and any correlation of the off-diagonal elements between GRMs will induce a bias. Therefore, we can show that the expected estimates for the three regressors under HE when performing a simple linear regression are:

$$\begin{aligned}\mathbb{E}(\hat{h}^2_{SNP}) &= h^2_{SNP} + \delta^2_{SNP} r_{\Theta_{A_{ij}}\Theta_{D_{ij}}} \sqrt{\frac{Var(\Theta_{D_{ij}})}{Var(\Theta_{A_{ij}})}} + \eta^2_{SNP} r_{\Theta_{A_{ij}}\Theta_{AA_{ij}}} \sqrt{\frac{Var(\Theta_{AA_{ij}})}{Var(\Theta_{A_{ij}})}} \\ &= h^2_{SNP} + \delta^2_{SNP} r_{\Theta_{A_{ij}}\Theta_{D_{ij}}} \sqrt{\frac{1}{2}} + \eta^2_{SNP} r_{\Theta_{A_{ij}}\Theta_{AA_{ij}}} \sqrt{\frac{2}{M}}\end{aligned}$$

$$\begin{aligned}\mathbb{E}(\hat{\delta}^2_{SNP}) &= \delta^2_{SNP} + h^2_{SNP} r_{\Theta_{D_{ij}}\Theta_{A_{ij}}} \sqrt{\frac{Var(\Theta_{A_{ij}})}{Var(\Theta_{D_{ij}})}} + \eta^2_{SNP} r_{\Theta_{D_{ij}}\Theta_{AA_{ij}}} \sqrt{\frac{Var(\Theta_{AA_{ij}})}{Var(\Theta_{D_{ij}})}} \\ &= \delta^2_{SNP} + h^2_{SNP} r_{\Theta_{D_{ij}}\Theta_{A_{ij}}} \sqrt{2} + \eta^2_{SNP} r_{\Theta_{D_{ij}}\Theta_{AA_{ij}}} \sqrt{\frac{4}{M}}\end{aligned}$$

$$\begin{aligned}\mathbb{E}(\hat{\eta}^2_{SNP}) &= \eta^2_{SNP} + h^2_{SNP} r_{\Theta_{AA_{ij}}\Theta_{A_{ij}}} \sqrt{\frac{Var(\Theta_{A_{ij}})}{Var(\Theta_{AA_{ij}})}} + \delta^2_{SNP} r_{\Theta_{AA_{ij}}\Theta_{D_{ij}}} \sqrt{\frac{Var(\Theta_{D_{ij}})}{Var(\Theta_{AA_{ij}})}} \\ &= \eta^2_{SNP} + h^2_{SNP} r_{\Theta_{AA_{ij}}\Theta_{A_{ij}}} \sqrt{\frac{M}{2}} + \delta^2_{SNP} r_{\Theta_{AA_{ij}}\Theta_{D_{ij}}} \sqrt{\frac{M}{4}}\end{aligned}$$

with $r_{\Theta_{A_{ij}}\Theta_{D_{ij}}}$ the correlation of the off-diagonal elements of the additive and dominance GRM and so on. The variance of the off-diagonal elements of $\Theta_A$ is $Var(\Theta_{A_{ij}}) = \frac{1}{M}$ with M the effective number of markers, for $\Theta_D$ we have $Var(\Theta_{D_{ij}}) \simeq \frac{2}{M}$ and $Var(\Theta_{AA_{ij}}) \simeq \frac{2}{M^2}$ (**Supplementary Note 2**). Having a closer look to these expectations reveals that the bias on $\eta^2_{SNP}$ estimates due to additive and dominance variance are proportional to $\sqrt{M}$

whereas the collinearity effect of additive-by-additive variance on the additive and dominance estimates is inversely proportional to $\sqrt{M}$.

Under the parametrization of our model and assuming HWE, linkage equilibrium between markers and that we know the true population allele frequencies, the genotype variable $x_{A(i)}$ and $x_{D(i)}$ are expected to be independent with $Cov(x_{A(i)}, x_{D(i)}) = 0$, leading to $r_{\Theta_{A_{ij}}\Theta_{D_{ij}}} = 0$. Moreover, since we assume $\Theta_{AA_{ij}} \sim \mathcal{N}(0, Var(\Theta_{A_{ij}}))$, we also expect $r_{\Theta_{A_{ij}}\Theta_{AA_{ij}}} = 0$, however, we do not have a theoretical expectation for $r_{\Theta_{D_{ij}}\Theta_{AA_{ij}}}$.

In practice, the true population allele frequencies are unknown and estimated from a particular sample. Consequently, the use of estimated allele frequencies to compute the different GRMs can results in a correlation structure between the off-diagonal elements of the GRMs, leading to a confounding effect. If this effect is expected to be very small on $\hat{h}^2_{SNP}$ and $\hat{\delta}^2_{SNP}$ regarding their expectations in a single regression analysis, it can result in a significant bias on $\hat{\eta}^2_{SNP}$ when using a simple regression with $\Theta_{AA}$ only.

We were able to reproduce the sampling effect on the bias of $\hat{\eta}^2_{SNP}$ using a simple example where we simulated a trait with $h^2_{SNP} = 0.5$ and $\delta^2_{SNP} = \eta^2_{SNP} = 0$ (following **Equation 5** of the main manuscript) for 1000 unrelated individuals genotyped at 1000 simulated unlinked SNPs with population allele frequencies $p = 0.5$. We compared $\hat{\eta}^2_{SNP}$ under HE regression when fitting both $\Theta_A$ and $\Theta_{AA}$, or $\Theta_{AA}$ only using 1000 replicates. We obtained unbiased estimates of $\eta^2_{SNP}$ when both $\Theta_A$ and $\Theta_{AA}$ were fitted in the model, whereas a negative bias was observed when using $\Theta_{AA}$ only in the model (**Supplementary Figure 9**). However, when the true population allele frequencies were used to compute the different GRMs, this bias disappeared and unbiased estimates of $\eta^2_{SNP}$ were obtained both under multiple and single HE regression analysis.

We computed the average correlation of the off-diagonal elements of the GRMs across the 1000 replicates. If we observed that $r_{\Theta_{A_{ij}}\Theta_{D_{ij}}}$ decreased from -1.00e-3 to -1.50e-7, and $r_{\Theta_{A_{ij}}\Theta_{AA_{ij}}}$ from -4.47e-2 to 1.27e-4 when using the true population allele frequencies instead of the estimated ones, we did not observe any effect on $r_{\Theta_{D_{ij}}\Theta_{AA_{ij}}}$ (2.25e-2 and 2.24e-02 when using estimated or true allele frequencies respectively).

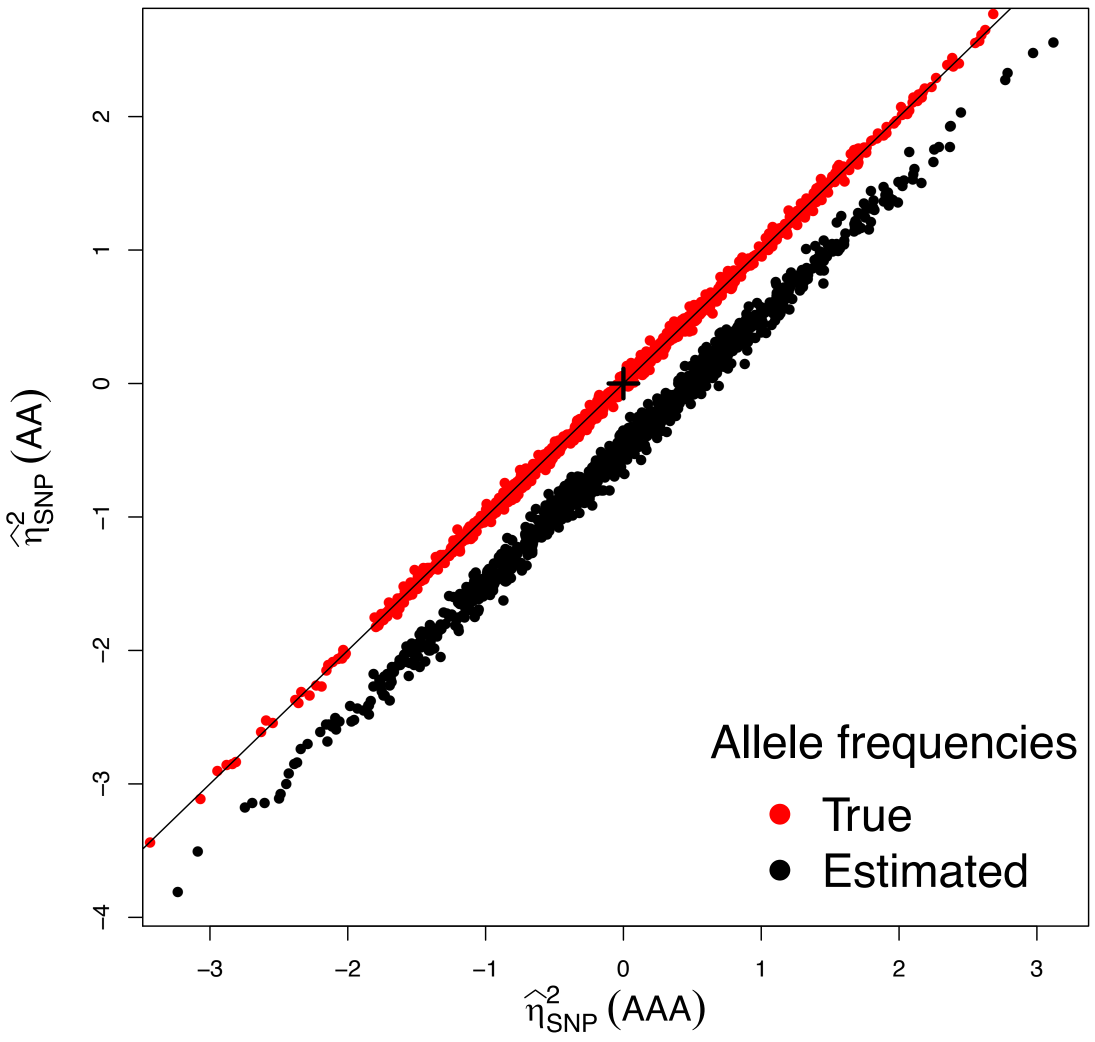


**Supplementary Figure 9: Single-replicate estimates from unlinked markers simulations using true or estimated allele frequencies to standardize the genotypes.** We compared HE estimates for 1000 replicates of simulations of 1000 unrelated individuals *genotyped at* 1000 simulated unlinked markers with population allele frequencies p = 0.5. We simulated phenotypes using all the markers as causal SNPs and with additive variance $h^2_{SNP} = 0.5$ and $\delta^2_{SNP} = \eta^2_{SNP} = 0$, following **Equation (5)** of the main manuscript, using either the true (red dots) or estimated (black dots) allele frequencies from the sampled individuals. We compared estimates of $\eta^2_{SNP}$ from

HE regression when $\Theta_A$ and $\Theta_{AA}$ are fitted together (AAA) or $\Theta_{AA}$ only (AA). The simulated value is depicted by a black cross and the black dashed line depicts the Y=X line.

**Supplementary Note 2**

**Sampling variance of the estimation of additive-by-additive variance from a sample of unrelated individuals**

**Supplementary Table 6: Summary of main notations.**

| Notation | Description |
|---|---|
| G=GRM | The genomic relationship matrix estimated from L markers |
| $\mathrm{Var}(G_{ij}) = \frac{1}{M}$ | The variance of the off-diagonal elements of G. M is the effective number of independent markers. If markers are unlinked, then M is simply the total number of markers L used to estimate G |
| $G \odot G$ | The Hadamard product of G with itself. This is the matrix used to estimate additive-by-additive variance |
| $h_{SNP}^2$ | Proportion of phenotypic variance explained by G |
| $\delta_{SNP}^2$ | Proportion of phenotypic variance explained by the dominance GRM |
| $\eta_{SNP}^2$ | Proportion of phenotypic variance explained by $G \odot G$ |

We aimed at deriving the approximate sampling variance of an estimate of $\eta_{\mathrm{SNP}}^2$, which can then be used in power calculations. A summary of the main notation use in this **Supplementary Note 2** are provided in the **Supplementary Table 3**.

Assuming that $G_{ii} \sim \mathcal{N}(1, Var(G_{ii}))$, $G_{ij} \sim \mathcal{N}\left(0, Var(G_{ij})\right)$ and $Var(G_{ij}) = \frac{1}{M}$, with M the effective number of markers, i and j are individuals. $Var(G_{ii})$ depends on the choice of markers and on how the diagonal elements are calculated[1].

Visscher et al.[2] derived the sampling variance of $\hat{h}^2_{SNP}$ as $\frac{2}{N^2 Var(G_{ij})} = \frac{2M}{N^2}$, where N is the total sample size. This derivation was for HE regression (using the phenotypic cross-products) but was found to be accurate enough for REML estimates too. Visscher & Goddard[3] showed that for REML estimation a general approximation of the sampling variance of $\hat{h}^2_{SNP}$ is $\frac{2}{NVar(\lambda_G)}$ with $\lambda_G$ the eigenvalues of matrix G. They also showed that in the case of G estimated from markers, $Var(\lambda_G) \simeq NVar(G_{ij})$, hence $var(\hat{h}^2_{SNP}) \simeq \frac{2}{[N^2 Var(G_{ij})]} = \frac{2M}{N^2}$, which is the same as the derivation from HE regression.

### Sampling variance of $\hat{\eta}^2_{SNP}$ for HE regression

For HE regression using the additive-by-additive GRM (G ⊙ G), defined as the square of the additive GRM G, it is possible to derive the expected variance of the off-diagonal elements. From the normality assumption, we have:

$$\begin{aligned} G_{ij} &\sim \mathcal{N}(0, \sigma^2) \\ &\sim \sigma\mathcal{N}(0,1) \end{aligned}$$

then we have:

$$\begin{aligned}(\mathrm{G} \odot \mathrm{G})_{ij} &\sim G_{ij}^2 \sim \sigma^2 \chi_1^2 \\ &\sim Var\big(G_{ij}\big)\chi_1^2 \\ &\sim \frac{\chi_1^2}{M}\end{aligned}$$

Hence:

$$Var\big((\mathrm{G} \odot \mathrm{G})_{ij}\big) = Var(G_{ij})^2 Var(\chi_1^2), \text{ with } Var(\chi_1^2) = 2$$

$$= 2Var(G_{ij})^2 \quad (1)$$

$$= \frac{2}{M^2} \quad (2)$$

Therefore, for HE regression, we have $Var\big(\hat{\eta}^2_{SNP_{HE}}\big) \simeq \frac{1}{N^2 Var(G_{ij})^2} = \left[\frac{M}{N}\right]^2$. As it has been shown empirically that $Var\big(G_{ij}\big) \simeq 2e-5$ (when computed from genome-wide data) for humans, we expect $Var((\mathrm{G} \odot \mathrm{G})_{ij}) \simeq 8e-10$. It is worth emphasizing that Equation 1 can be used for a particular sample after the GRM has been observed. However, Equation 2 can be used for power calculation before a particular GRM has been observed. Indeed, assuming a sample of N unrelated individuals genotyped at unlinked markers, then M simply becomes the number of markers L.

### Sampling variance of $\hat{\eta}^2_{SNP}$ for REML

We use the result from Visscher & Goddard[3] and applied it to the case where REML is used with $\mathrm{G} \odot \mathrm{G}$ only. Using this result, we expect $Var\big(\hat{\eta}^2_{SNP_{REML}}\big)$to be $\frac{2}{NVar(\lambda_{(\mathrm{G}\odot\mathrm{G})})}$. Then we derive the mean eigenvalue $\mathbb{E}\big(\lambda_{(\mathrm{G}\odot\mathrm{G})}\big)$:

$$
\begin{aligned}
\mathbb{E}\big(\lambda_{(\mathrm{G}\odot\mathrm{G})}\big) &= \frac{Tr(\mathrm{G}\odot\mathrm{G})}{N} \\
&= \frac{\sum_{i;j}^{N} G_{ij}^{2}}{N} \\
&= 1 + Var(G_{ii})
\end{aligned}
$$

To derive the variance of $\lambda_{(\mathrm{G}\odot\mathrm{G})}$, we use the fact that $\sum_{i;j}^{L}(\mathrm{G}\odot\mathrm{G})_{ij}^{2} = \sum\lambda_{(\mathrm{G}\odot\mathrm{G})}^{2}$, i.e the sum of squares of all elements in G ⊙ G is equal to the sum of squares of its eigenvalues. Then, from normality assumptions, we have:

$$
\begin{aligned}
\sum_{i;j}^{L}(\mathrm{G}\odot\mathrm{G})_{ij}^{2} &= N(N-1)\mathbb{E}\big(G_{ij}^{4}\big) + N\mathbb{E}(G_{ii}^{4}) \\
&= 3N(N-1)Var(G_{ij})^{2} + N + 6NVar(G_{ii}) + 3NVar(G_{ii})^{2}
\end{aligned}
$$

Therefore,

$$
\begin{aligned}
Var\big(\lambda_{(\mathrm{G}\odot\mathrm{G})}\big) &= \frac{\sum\lambda_{(\mathrm{G}\odot\mathrm{G})}^{2}}{N} - \mathbb{E}\big(\lambda_{(\mathrm{G}\odot\mathrm{G})}\big)^{2} \\
&= \frac{1}{N}\sum_{i;j}^{N}(\mathrm{G}\odot\mathrm{G})_{ij}^{2} - (1 + Var(G_{ii}))^{2} \\
&= 4Var(G_{ii}) + 2Var(G_{ii})^{2} + 3(N-1)Var\big(G_{ij}\big)^{2} \qquad (3) \\
&= 4Var(G_{ii}) + 2Var(G_{ii})^{2} + \frac{3(N-1)}{M^{2}}
\end{aligned}
$$

Finally,

$$
\begin{aligned}
Var\big(\hat{\eta}_{SNP_{REML}}^{2}\big) &\simeq \frac{2}{NVar\big(\lambda_{\mathrm{G}\odot\mathrm{G}}\big)} \\
&\simeq \frac{2}{N(4Var(G_{ii}) + 2Var(G_{ii})^{2} + 3(N-1)Var(G_{ij})^{2})} \qquad (4)
\end{aligned}
$$

For large sample size, and since $Var(G_{ii}) \sim O\left(Var\big(G_{ij}\big)\right) = O\left(\frac{1}{M}\right)$, we can simplify the equations to:

$$
\begin{aligned}
Var\left(\lambda_{(\mathrm{G}\odot\mathrm{G})}\right) &\simeq 4Var(G_{ii}) + 3NVar\left(G_{ij}\right)^2 \\
&\simeq 4Var(G_{ii}) + \frac{3N}{M^2} \\
Var\left(\hat{\eta}^2_{SNP_{REML}}\right) &\simeq \frac{(M/N)^2}{1.5 + 2M^2 Var(G_{ii})/N}
\end{aligned}
$$

The numerator $(M/N)^2$ is the approximate sampling variance for HE regression. Therefore,

$$
\begin{aligned}
\frac{Var\left(\hat{\eta}^2_{SNP_{HE}}\right)}{Var\left(\hat{\eta}^2_{SNP_{REML}}\right)} &\simeq 1.5 + 2M^2 \frac{Var(G_{ii})}{N} \\
&\simeq 1.5 \text{ for } N \to \infty
\end{aligned}
$$

As for equation 1 with HE, equation 3 can be used for REML power calculation when the GRM is already observed. However, to have an approximation of the sampling variance of $\hat{\eta}^2_{SNP}$ before computing a GRM, we need to approximate $Var\left(G_{ij}\right)$ and $Var(G_{ii})$. Again, if we consider a sample of N unrelated individuals genotyped at unlinked markers, then $M = L$ the total number of markers. Moreover, Yang et al.[1] derived the variance of the diagonal element of G at one marker k as:

$$
Var(G_{iik}) = \frac{1}{2p_k(1-p_k)} - 1
$$

with $p_k$ the allele frequency at locus k. Then, each diagonal element of the GRM is compute following a common-sense weighting scheme with $G_{ii} = \frac{1}{M}\sum_j^M G_{iik}$. Hence:

$$
Var(G_{ii}) = \frac{1}{M}\sum_k^M V\,ar(G_{iik})
$$

As noticed previously, $Var(G_{ii})$ depends both on the way to compute the diagonal elements and on the choice of markers, more precisely the distribution of allele frequencies. Then, assuming we use a minimum MAF of 1%, and assuming a uniform distribution of allele frequencies on [0.01; 0.99], we can approximate:

$$\begin{aligned} Var(G_{ii}) &= \frac{1}{M}\sum_{k}^{M} Var(G_{iik}) \\ &= \frac{1}{M}\int_{0.01}^{0.99} \frac{1}{2p(1-p)} - 1\,dp \\ &\simeq 3.6 \times \frac{1}{M} \\ &\simeq 3.6 \times Var(G_{ij}) \end{aligned}$$

Hence,

$$Var(\lambda_{G\odot G}) \simeq 14Var(G_{ij}) + 3NVar(G_{ij})^2$$

$$Var(\lambda_{G\odot G}) \simeq \frac{14}{M} + \frac{3N}{M^2} \quad (5)$$

And,

$$Var(\hat{\eta}^2_{SNP_{REML}}) \simeq \frac{2}{N\left(\frac{14}{M} + \frac{3N}{M^2}\right)} \quad (6)$$

$$\simeq \frac{2}{3} Var(\hat{\eta}^2_{SNP_{HE}}) \quad (7)$$

We used 100 simulations of 500 unrelated individuals genotyped at 10,000 simulated unlinked SNPs with uniform allele frequencies distribution on [0.01;0.99]. The **Supplementary Figure 10** shows that estimates of $Var(\lambda_{(G\odot G)})$ of the correspondings additive-by-additive GRM are highly accurate when using the equation 4. However, the **Supplementary Figure 11** shows that approximation from equation 6 lead to biased estimates of $Var(\hat{\eta}^2_{SNP_{REML}})$, somewhat small regarding the average estimate ($\simeq$ 2.82) compare to the average true value ($\simeq$ 2.69).

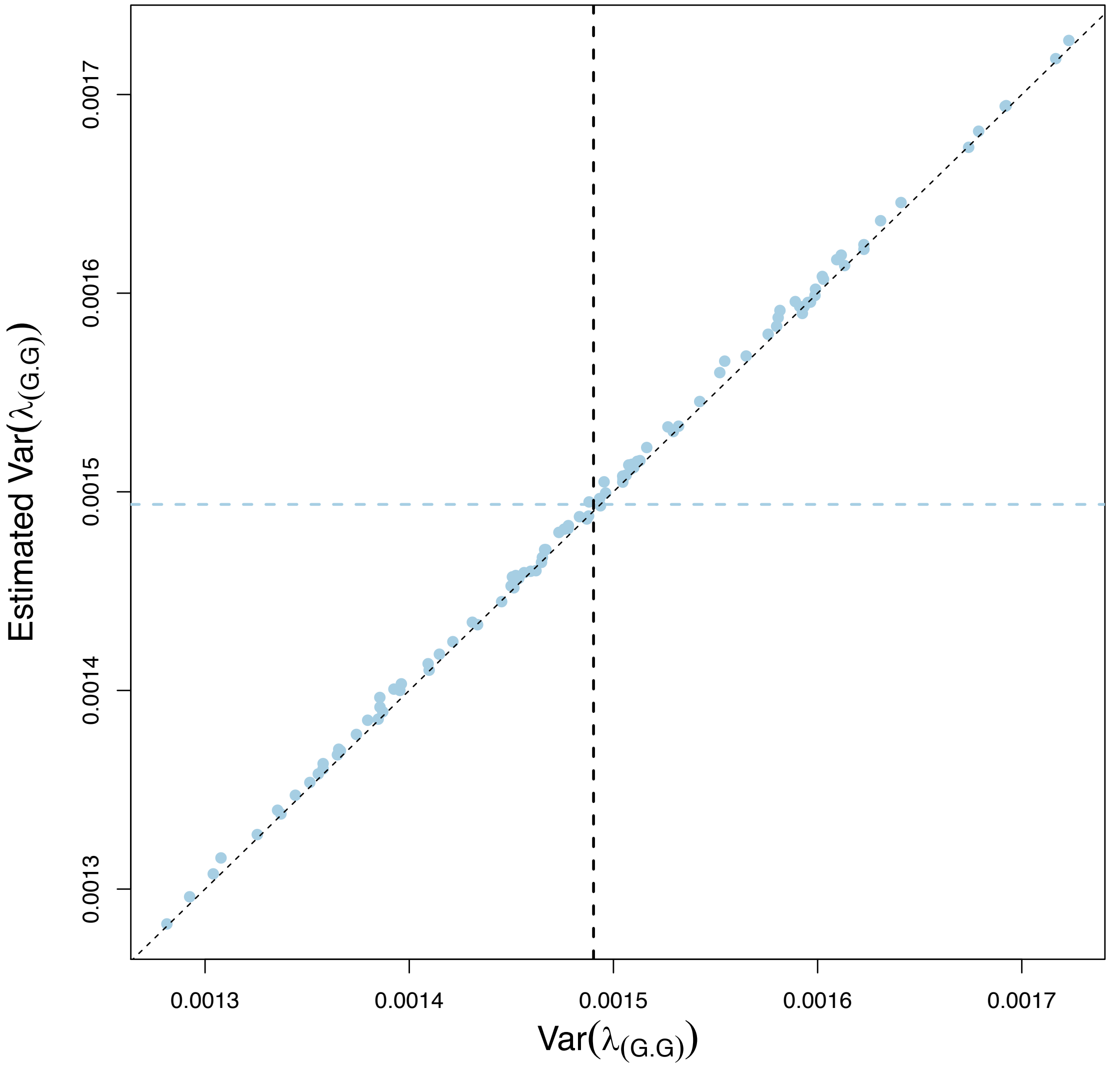


**Supplementary Figure 10: Plot of the simulated $Var(\lambda_{G\odot G})$ against the estimates from Equation (3) (blue dots).** We performed 100 replicates where Additive GRMs have been computed randomly for 500 unrelated individuals genotyped at 10,000 simulated unlinked SNPs with uniform distribution of allele frequencies on [0.01;0.99]. The vertical black dashed line depicts the average simulated $Var(\lambda_{G\odot G})$ whereas the horizontal blue dashed line depicts the average estimated $Var(\lambda_{G\odot G})$.

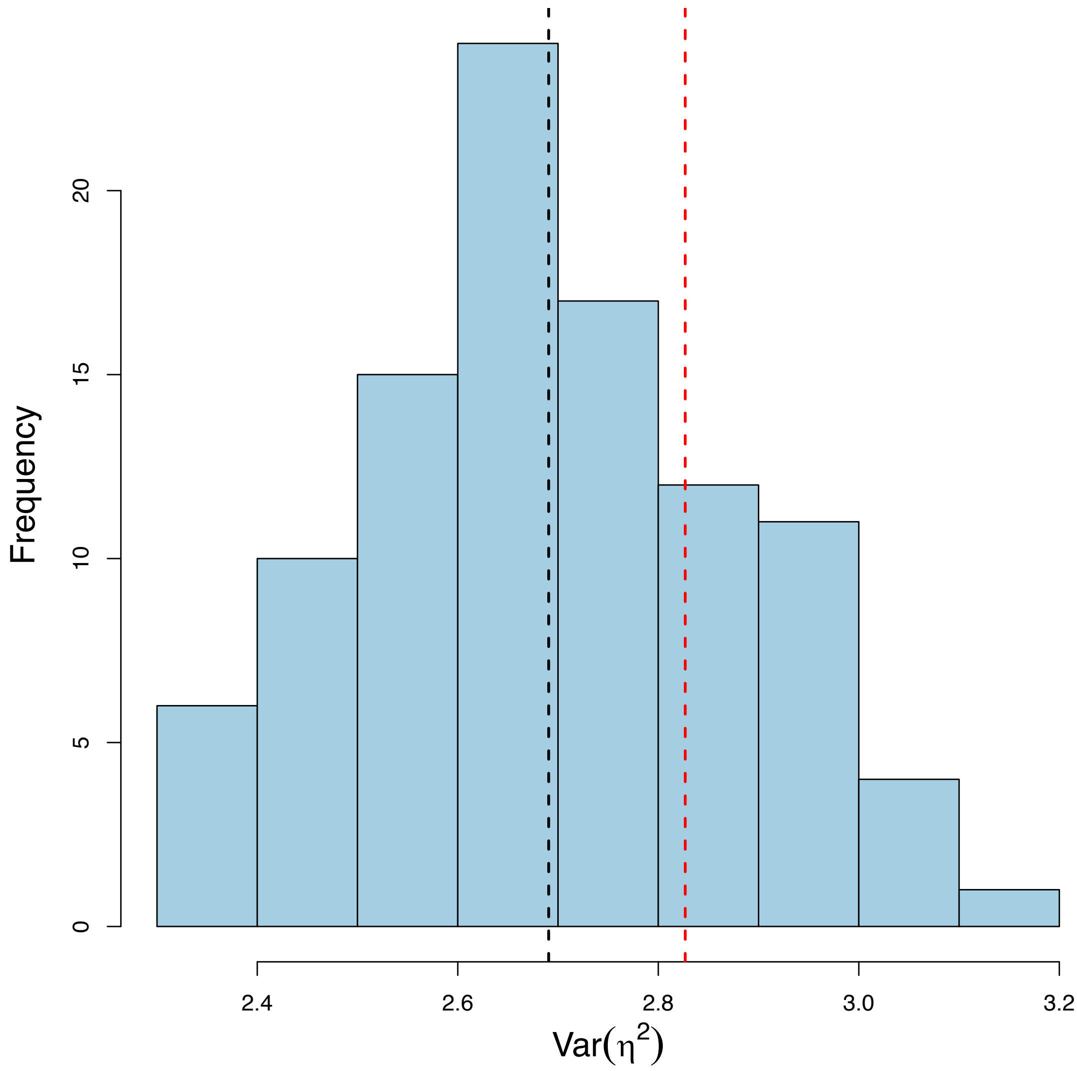


**Supplementary Figure 11: Distribution of the estimated $Var(\hat{\eta}^2_{SNP_{REML}})$ from simulations using Equation (4).** We performed 100 replicates where Additive GRMs have been computed randomly for 500 unrelated individuals genotyped at 10,000 simulated unlinked SNPs with uniform distribution of allele frequencies on [0.01;0.99]. The average estimate is depicted with the black dashed line whereas the red dashed line shows the approximation of $Var(\hat{\eta}^2_{SNP_{REML}})$ using **Equation (6)**.

**Supplementary Note 3**

**Small effect sizes lead to additivity.**

GWAS results are consistent with highly polygenetic architectures, which implies that at any one locus the average effect sizes are very small, relative to the mean or standard deviation of the trait, irrespective of the scale of measurement. This by itself predicts that interactions within loci (dominance) or between loci (epistasis) are likely to be small and that (therefore) dominance or epistatic variance is small relative to non-additive variance[4].

Even highly nonlinear biological systems can lead to mostly additive genetic variance. Keightley[5] used metabolic controls theory to model the relationship between enzyme activity and flux, which is known to have a non-linear association. He showed that unless effect sizes of mutations on enzyme activity are large, most genetic variance in the phenotype flux is additive.

Here we give a simple but general derivation why dominance effects at individual loci are likely to be much smaller than additive effects. The same reasoning also applies to epistatic interactions.

Let $\mathbb{E}(y|x) = \mu + \beta x$, with m the mean, $x$ = 0, 1, 2 and $\beta$ the average effect sizes on the $y$-scale. Now consider any continuous functional relationship $f(y)$, which can be highly non-linear. Using a 2nd order Taylor series approximation gives,

$$f(\mu + 0 \times \beta) = f(\mu)$$

$$f(\mu + \beta) \simeq f(\mu) + \beta f'(\mu) + \frac{1}{2}\beta^2 f''(\mu)$$

$$f(\mu + 2\beta) \simeq f(\mu) + 2\beta f'(\mu) + 2\beta^2 f''(\mu)$$

with $f'(\mu)$ and $f''(\mu)$ the first and second derivative of $f(y)$ with respect to $y$, evaluated at $y = \mu$. The additive and dominance coefficient of $f(y)$ are then, approximately,

$$\mathrm{a}_f \simeq \frac{1}{2}[f(\mu) + 2\beta f'(\mu) + 2\beta^2 f'(\mu) - f(\mu)] = \beta f'(\mu) + \beta^2 f''(\mu)$$

and,

$$\mathrm{d}_f \simeq \left[f(\mu) + \beta f'(\mu) + \frac{1}{2}\beta^2 f''(\mu)\right] - \frac{1}{2}[2f(\mu) + 2\beta f'(\mu) + 2\beta^2 f''(\mu)]$$

$$\simeq -\frac{1}{2}\beta^2 f''(\mu)$$

Therefore, the ratio of dominance to additive coefficient on the $f(y)$ scale is,

$$\frac{\mathrm{d}_f}{\mathrm{a}_f} \simeq -\frac{1}{2}\frac{\beta f''(\mu)}{f'(\mu)}$$

This ratio tends to 0 when $\beta$ → 0 and/or when $f''(\mu)/\, f'(\mu)$ → 0.

As an example, let $\boldsymbol{f(y) = y^k}$, which can be very non-linear. Then $\boldsymbol{f'^{(y)} = ky^{k-1}}$, $\boldsymbol{f'^{(y)} = k(k-1)y^{k-2}}$, and $\frac{\mathbf{d}_f}{\mathbf{a}_f} \simeq -\frac{1}{2}\frac{\boldsymbol{\beta(k-1)}}{\boldsymbol{\mu}}$, which is small when $\boldsymbol{\beta/\mu}$ is small. Note that when $\frac{\mathbf{d}_f}{\mathbf{a}_f}$ is small, then $\frac{\sigma_D^2(f(y))}{\sigma_A^2(f(y))}$ is even smaller.
